# MAIT cells protect against sterile lung injury

**DOI:** 10.1101/2024.01.08.574631

**Authors:** Xiawei Zhang, Shuailin Li, Wojciech Lason, Maria Greco, Paul Klenerman, Timothy SC Hinks

## Abstract

Mucosal-associated invariant T (MAIT) cells, the most abundant unconventional T cells in the lung, have been recently linked to tissue protection and repair. Their role, especially in sterile lung injury, is unknown. Using single cell RNA sequencing (scRNA-seq), spectral analysis and adoptive transfer in a bleomycin-induced sterile lung injury, we found that bleomycin activates murine pulmonary MAIT cells and induces an accompanying tissue repair programme, associated with a protective role against bleomycin-induced lung injury. MAIT cells drive the accumulation of type 1 conventional dendritic cells (cDC1), limiting tissue damage in a DNGR-1 dependent manner. Human scRNA-seq data revealed that MAIT cells were activated, with increased cDC populations in idiopathic pulmonary fibrosis patients. Thus, MAIT cells enhance defence against sterile lung injury by fostering cDC1-driven anti-fibrotic pathways.

## INTRODUCTION

Mucosal-associated invariant T (MAIT) cells are innate-like T cells that recognise small molecule derivatives of riboflavin synthesis ^1^ such as 5-(2-oxopropylideneamino)-6-d-ribitylaminouracil (5-OP-RU) presented on the major histocompatibility complex (MHC)-related protein-1 (MR1) ^2, 3^. MAIT cells have potential for multiple diverse functions in the host response to a wide variety of bacterial, viral and fungal pathogens and in promoting tissue repair ^4–7^. MAIT cells are particularly abundant in the lung, comprising up to 10% of all pulmonary T cells in a healthy human ^8, 9^. They are characterised by expression of a semi-invariant T-cell receptor (TCR)-α chain: typically Vα7.2–Jα33/12/20 in humans and Vα19– Jα33 in mice^2, 3^. These features imply an essential role of MAIT cells in pulmonary mucosal immunology, particularly during the initial stages of an immune response, yet our understanding of the full repertoire of MAIT cell functions remains incomplete, particularly in the context of tissue injury and repair. Increasing evidence implicates MAIT cells in bridging innate and adaptive immunity, an important role being the recruitment of other immune cells, particularly dendritic cells (DC) and monocytes during TCR-dependent ^10–12^ and cytokine-dependent activation ^13, 14^.

Strategically located within the airway epithelium and interstitium, pulmonary DCs bridge the external and internal environments ^15, 16^. The lung features two distinct conventional dendritic cell subsets (cDC, MHCII^+^ CD11c^+^): CD103^+^ type 1 cDC (CD103^+^ CD11b^lo/–^ XCR1^+^ DNGR-1^+^ SIRP-α^−^ CX3CR1^−^ F4/80^−^, cDC1) and CD11b^+^ type 2 cDC (CD11b^hi^ CD103^−^SIRP-α^+^ CX3CR1^+^ F4/80^+^, cDC2) ^17^. cDC1 specialize in cross-presenting antigens to CD8^+^ T cells, promoting Th1 cells. In contrast, cDC2 excel at stimulating CD4^+^ T cell responses, mainly Th2 or Th17 cells. Lung cDCs originate from common dendritic cell precursors (CDPs) in the bone marrow. These CDPs mature into pre-dendritic cells (pre-DCs), which migrate to the lungs through the bloodstream and differentiate into either cDC1 or cDC2 subsets guided by local signals and specific transcription factors ^18–20^.

DCs play a regulatory role in pulmonary fibrosis, accumulating in the lungs in idiopathic pulmonary fibrosis (IPF) patients^21–23^ and in bleomycin mouse models^24, 25^, whilst diminishing in the circulation^26^. Pulmonary cDC1s increase with bleomycin treatment, but are reduced with transforming growth factor (TGF)-β inhibition, suggesting anti-inflammatory and anti-fibrotic roles in pulmonary fibrosis^25^. Moreover, increased fibrosis severity and impaired lung function were seen in DC-deficient mice, but mitigated when DC counts were boosted^27^, though their protective mechanism is still unclear.

In this study, we aim for the first time to define the role of MAIT cells in sterile lung injury and to investigate the underlying mechanisms. We employed a model of lung injury using bleomycin: a potent chemotherapeutic agent, with a well-characterised side effect profile of acute lung injury, followed by a chronic phase with pathological hallmarks of human IPF ^28^. We have shown for the first-time that MAIT cells accumulate and are activated upon sterile injury, in a cytokine-dependent manner, and we have discovered an *in vivo* mechanism by which pulmonary MAIT cells make an important contribution to protection against sterile lung tissue damage in mice.

## RESULTS

### Activated MAIT cells accumulate in the lung upon bleomycin treatment in a cytokine-dependent manner

Our initial objective was to establish whether sterile lung injury could stimulate pulmonary MAIT cells *in vivo*. We administered bleomycin intratracheally to C57BL/6 (wild type, WT) mice, to precipitate acute, sterile lung inflammation, followed by a tissue repair phase and subsequent fibrosis over a fortnight ^28^. We detected an early surge of pulmonary MAIT cells (characterised as CD45.2^+^ TCRβ^+^ CD19^−^ MR1-5-OP-RU tetramer^+^ cells, Supplementary Fig. 1A), peaking approximately three days post-challenge. By comparison, the peak number of non-MAIT αβ T cells occurred later at day 7 post-challenge (Fig. 1A and Supplementary Fig. 2, G to H). On both day 3 and 5 post-challenge, the fold change in MAIT cell frequency was significantly higher than that in non-MAIT αβ T cells frequency (Fig. 1B and Supplementary Fig. 2, J to K). Moreover, pulmonary MAIT cell CD69 expression increased significantly at days 3 and 5 post challenge relative to unchallenged controls, and CD69 expression was significantly higher in MAIT cells than that in non-MAIT αβ T cells (Fig. 1C, Supplementary Fig. 2, I and L).

**Fig. 1:**
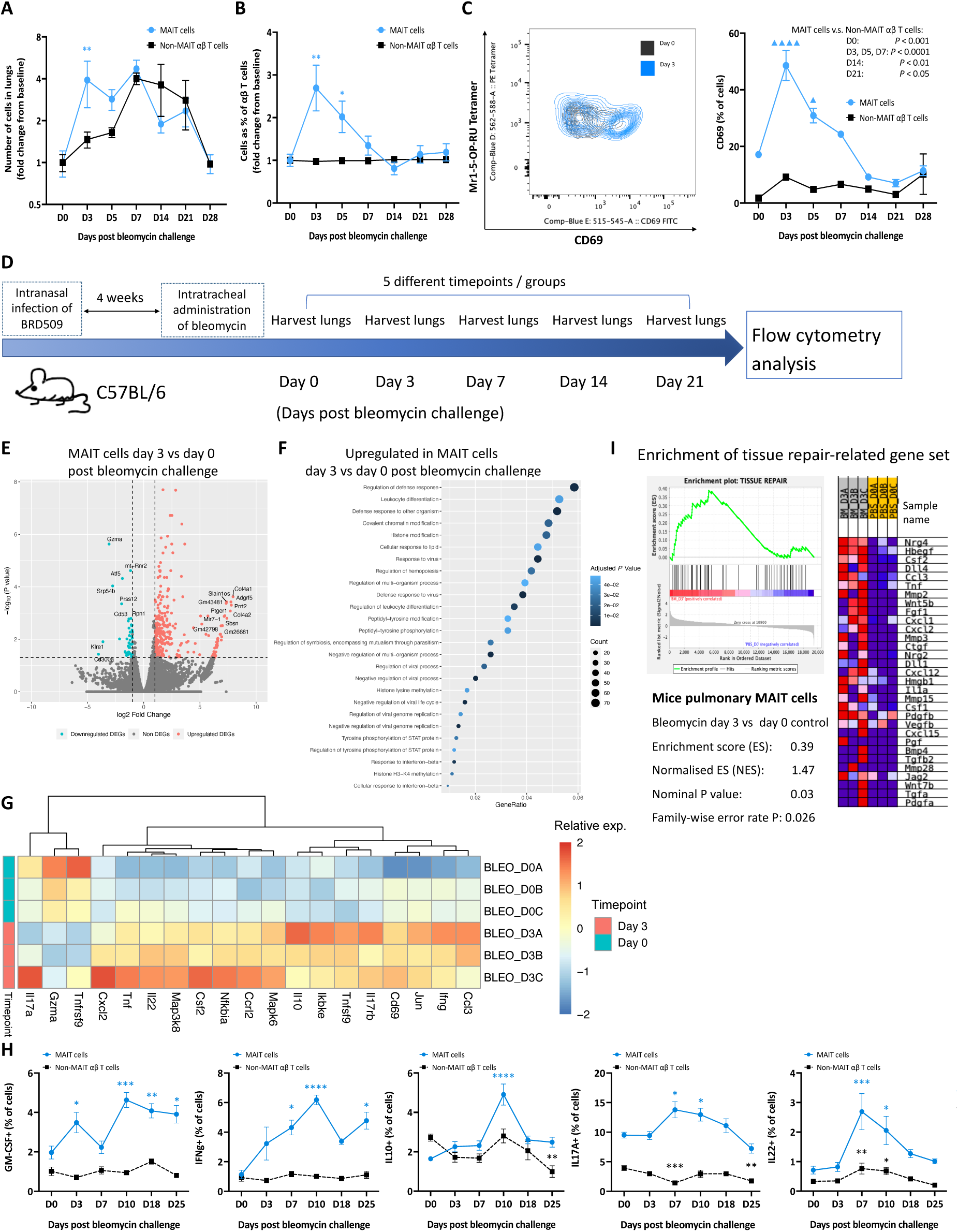
*In vivo* bleomycin challenge induces cytokine-modulated accumulation and activation of pulmonary MAIT cells, triggering their tissue repair programme. (**A**) Fold change in the absolute MAIT cell and non-MAIT αβ T cell number in the lungs of naïve mice post-bleomycin challenge, relative to unchallenged PBS controls (Day 0). Comparisons between MAIT cell subsets and non-MAIT αβ T cell subsets at individual time-points, were conducted using unpaired t tests or Mann-Whitney tests, \**P* < 0.05, \*\**P* < 0.01, \*\*\**P* < 0.001, \*\*\*\**P* < 0.0001. (**B**) Fold change in MAIT cell and non-MAIT αβ T cell frequency as the percentage of total pulmonary αβ T cells in the lungs of naïve mice relative to unchallenged controls. Comparisons between MAIT cell subsets and non-MAIT αβ T cell subsets at individual time-points, were conducted using unpaired t tests or Mann-Whitney tests, \**P* < 0.05, \*\**P* < 0.01, \*\*\**P* < 0.001, \*\*\*\**P* < 0.0001. (**C**) Proportion of pulmonary MAIT cells and non-MAIT αβ T cells expressing CD69 in naïve mice. Statistical comparisons across different timepoints post-bleomycin challenge and unchallenged PBS control were made using one-way ANOVA with Dunnett’s or Dunn’s multiple comparison tests,▴*P* < 0.05, ▴▴*P* < 0.01, ▴▴▴*P* < 0.001, ▴▴▴▴*P* < 0.0001. Comparisons between MAIT cell subsets and non-MAIT αβ T cell subsets at individual time-points, were conducted using unpaired t tests or Mann-Whitney tests. The data are presented as the mean ± SEM of a single experiment, with 3-5 mice in each group. (**D**) Schematic of protocol: Both WT and Mr1^−/−^ mice were infected with 10^6^ CFU *S. typhimurium* BRD509 to augment the MAIT cell population, followed by intratracheal bleomycin administration. Mouse lung samples were collected at days 0, 3, 7, 14 and 21 after the challenge. (**E**) Volcano plot of DEGs [log2 fold change (FC) > 1, adjusted *P* < 0.05] of mouse lung MAIT cells on day 3 post-bleomycin challenge compared with unchallenged PBS controls. The top 10 up and down-regulated genes are annotated. Horizontal line indicates *P* value threshold of 0.05. Vertical line indicates log2 fold change threshold of 1. (**F**) Top 25 significantly enriched (*P* < 0.05) pathways from Gene Ontology (GO) database (biological process) among upregulated DEGs in mouse lung MAIT cells at day 3 post-bleomycin challenge compared with unchallenged PBS controls. The colour intensity indicates the statistical significance of the enrichment, with the dot size representing the number of genes upregulated in each pathway. The x-axis illustrates the proportion of all DEGs included in each pathway (Gene Ratio). (**G**) Heatmap depicting expression of cytokines and chemokines in lung MAIT cells at days 0 (PBS control) and 3 post bleomycin challenge (red, highest expression; blue, lowest). (**H**) Proportion of pulmonary MAIT cells expressing GM-CSF, IFN-γ, IL-10, IL17A and IL22 expressed as percentage of MAIT and non-MAIT αβ T cells. Graphs show combined data (mean ± SEM) from two independent experiments with similar results, with 3– 5 mice per group in each replicate. Statistical tests compare each time-point with day 0 control in each subset by one-way ANOVA with Dunnett’s multiple comparisons test or Kruskal-Wallis with Dunn’s multiple comparisons test; \**P* < 0.05, \*\**P* < 0.01, \*\*\**P* < 0.001, \*\*\*\**P* < 0.0001. (**I**) GSEA for tissue repair gene signature ^35^. Heatmap showing the expression of leading-edge gene subsets in mice pulmonary MAIT cells at day 3 post-bleomycin challenge and unchallenged control (red indicates highest expression; blue indicates lowest).

In nature, early life microbial exposures are essential for the development of MAIT cell populations. When mice have been raised in a specific pathogen free environment, MAIT cells constitute less than 1% of total pulmonary αβ T cells ^29^ but can be up to 10% in healthy humans ^8, 30^. Therefore, we employed an established MAIT cell-enriched model by infecting WT C57BL/6 mice intranasally with 10^6^ CFU *Salmonella typhimurium* BRD509 four weeks prior to initial bleomycin challenge for clearer population delineation (Fig. 1D and Supplementary Fig. 1, B to D) ^31, 32^. This bacterial inoculum is rapidly cleared from the lungs but transiently provides the required combination of MAIT cell ligands and pathogen-associated molecular patterns necessary to produce rapid and lasting expansion of the MAIT cell population ^33^. While it was previously reported that post-*Salmonella* BRD509 infection MAIT cells adopt an effector memory phenotype, with relatively high baseline CD69 expression (Supplementary Fig. 1E) ^31^, we still observed rapid MAIT cell accumulation, peaking on day 3 post-challenge (Supplementary Fig. 2, A and B, M and N), along with activation upon bleomycin stimulation, evidenced by significantly increased CD69 expression on day 3 and 7, compared to unchallenged PBS controls (Supplementary Fig. 2, C and O). Conversely, no significant changes were observed in either the accumulation or activation of non-MAIT αβ T cells post-bleomycin challenge (Supplementary Fig. 2, A and B, P to R). Collectively, these results indicate MAIT cells accumulate and are activated early in the lungs following bleomycin-induced sterile injury of mice.

As MAIT cells can be activated in a TCR-independent manner by cytokines, including interleukin (IL)-12, -15, -18, and type I interferon (IFN), in antiviral responses ^34^, we examined MAIT cell responses post bleomycin challenge utilising mouse strains deficient in the two most important of these pathways, IL-18R or IFN-αR without a preliminary MAIT boost. Relative to WT C57BL/6 mice, we observed a marked impairment in pulmonary MAIT cell accumulation on day 3 post-bleomycin in both IL-18R and IFNαR-deficient mice (Supplementary Fig. 2, D and E, S and T, V and W). Conversely MAIT cell activation was significantly impeded solely in the absence of IFNαR (Supplementary Fig. 2, F, U and X). These findings suggest bleomycin-induced MAIT cell activation is predominantly cytokine driven, with IFN-αR playing a key role.

A transcriptional programme related to tissue repair, initially identified in Tc17 cells ^35^, has been observed in both human MAIT cells following 5-OP-RU stimulation and mice lung MAIT cells during acute *L. longbeachae* infection ^36^. Consequently, we assessed whether bleomycin could activate the MAIT cell tissue repair programme after bleomycin challenge in lungs of *S. typhi*-treated mice using bulk RNA-seq of flow-sorted pulmonary MAIT cells. The numbers of DEGs in bleomycin-challenged lung MAIT cells compared with unchallenged controls, were 425 (361 up, 64 down), 1230 (399 up, 831 down), 0, 24 (4 up, 20 down) and 131 (40 up, 91 down) genes at day 3, 7, 14, 21 and 28 post-bleomycin challenge, respectively (Supplementary Fig. 3A, for full list of DEGs see data S1). Intriguingly, the top 15 upregulated genes in pulmonary MAIT cells at day 3 post-bleomycin included tissue-damage related genes such as Col4α1, Col4α2, Ptger1, and Wwtr1 (Fig. 1E). The predominant GO pathways upregulated in MAIT cells at day 3 post-bleomycin challenge compared to those from unchallenged mice were associated with the regulation of the defence response, leukocyte differentiation, and response to viruses (Fig. 1F). We also observed a notable rise in the Cd69 gene expression and a modest upregulation of several inflammatory cytokines, such as Csf2, Ifn-γ, Tnf, Il17a, Il10 and Il22, in MAIT cells (Fig. 1G), and verified selected cytokines by flow cytometry (Fig. 1H and S3B).

We then conducted a gene set enrichment analysis (GSEA) to compare the TCR-triggered MAIT cell tissue repair gene signature ^36–38^, previously characterised in response to commensal bacteria in H2-M3-restricted RORγt^+^ CD8^+^ T cells in murine skin ^35^, with the gene expression profile of pulmonary MAIT cells following bleomycin exposure. This revealed significant enrichment of the tissue repair gene set in MAIT cells at day 3 post-bleomycin challenge (Fig. 1I). These findings suggested that *in vivo* bleomycin challenge activates the tissue repair programme of MAIT cells.

Current research indicates that MAIT cells are activated by MR1-TCR dependent or cytokine-dependent pathways; with the latter being key in antiviral response. Consequently, we further expanded our inquiry to investigate a different, albeit highly pertinent, condition of cytokine-dependent MAIT cell activation – viral infection. We performed bulk RNA-seq of MAIT cells isolated from the lungs of mice infected with 100 plaque forming units (PFU) of the mouse-adapted influenza virus strain A/Puerto Rico/8/34/1934 (PR8, H1N1). As expected, key upregulated genes included interferon-inducible genes, such as Ifi205 and Ifi44 (Supplementary Fig. 4, A and B, data S2). Upregulated GO terms included those related to defence responses against viruses and other organisms (Supplementary Fig. 4C). Intriguingly, our time-series analysis (Supplementary Fig. 4D) uncovered a tissue repair-oriented cluster – Cluster 2, characterised by enriched GO terms such as wound healing and regulation of cell morphogenesis (Supplementary Fig. 4, E and F). Importantly, the previously discussed tissue repair programme ^35^, was also activated in the MAIT cells at early timepoints post-infection (Supplementary Fig. 4, G to J), echoing the pattern we identified post-bleomycin, implying activation of this programme in both viral and sterile lung injury.

### MAIT cell-deficient mice show dysregulated pulmonary immune responses upon bleomycin challenge

We next sought to determine whether the recruitment and activation of MAIT cells in response to bleomycin have an impact on the phenotype. To this end, we assessed weight loss and tissue damage between WT and MAIT cell-deficient Mr1^−/−^ mice. WT and Mr1^−/−^ mice were subjected to 10^6^ CFU *S. typhimurium* BRD509 infection to expand the MAIT cell population followed by intratracheal bleomycin administration four weeks post-MAIT cell enrichment (Fig. 1D). Notably, Mr1^−/−^ mice exhibited more substantial weight loss (Fig. 2A), heightened tissue damage (Fig. 2B), and increased gene expression of Col1α1 and Col3α1 compared to WT mice (Fig. 2, C and D). Hydroxyproline levels in the lungs showed no significant difference between WT and Mr1^−/−^ mice (Supplementary Fig. 5A).

**Fig. 2:**
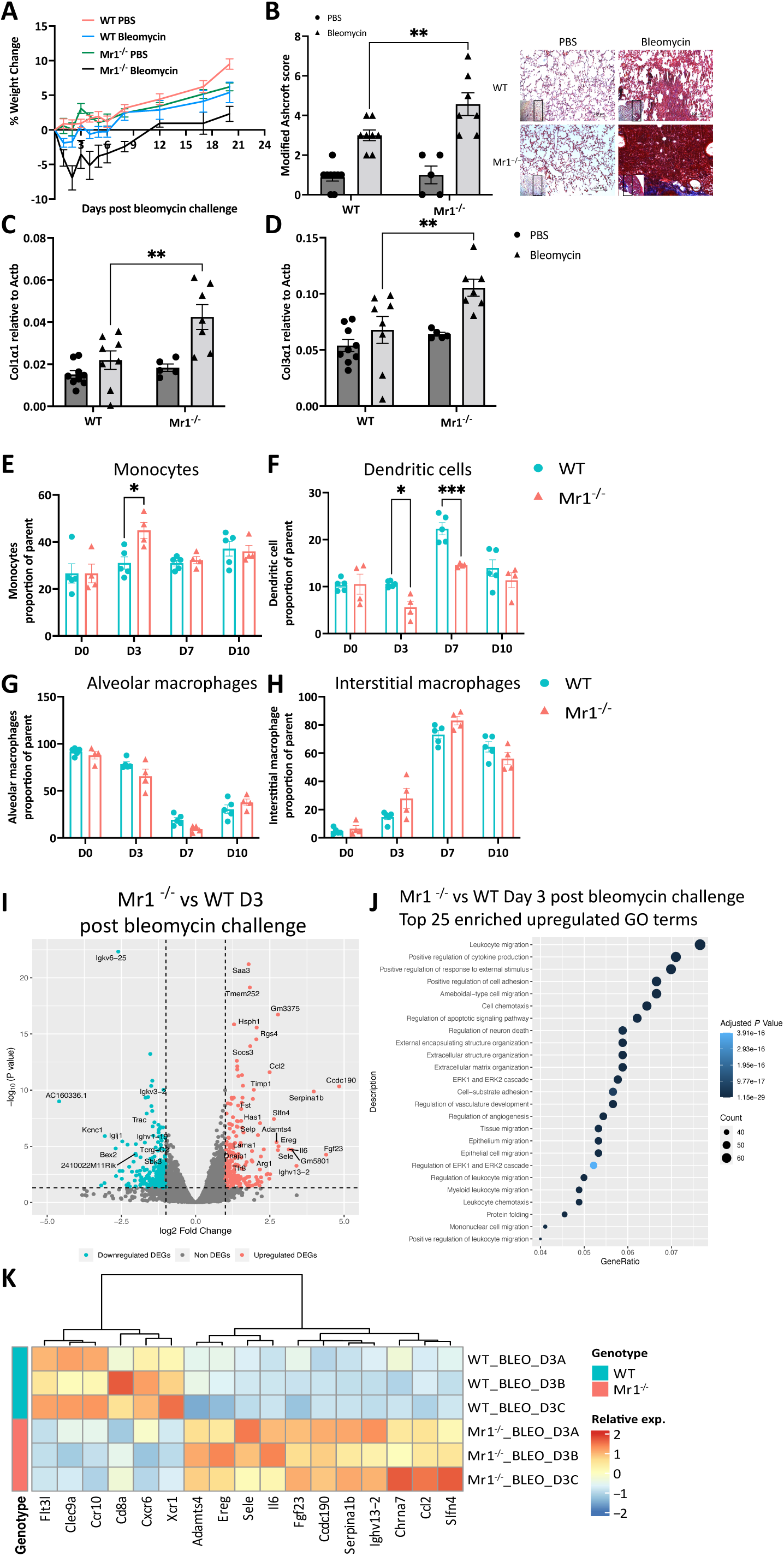
Dysregulated immune responses in the lungs of MAIT cell-deficient mice following bleomycin challenge. (**A**) Body weight loss expressed as a percentage of the weight before bleomycin challenge. (**B** and **C**) Modified Ashcroft score (**B**) and representative images (**C**) of lung slices of PBS or bleomycin-challenged WT and Mr1^−/−^ mice at day 21, stained with Masson’s trichrome. (**D**) Gene expression of Col1α1, and Col3α1 in lung homogenates of PBS or bleomycin-challenged WT and Mr1^−/−^ mice at day 21. Actb was used as a housekeeping gene. (**E** to **H**) Frequencies of monocytes (**E**), dendritic cells (**F**), alveolar macrophages (**G**) and interstitial macrophages (**H**) as percentages of parent in WT and Mr1^−/−^ mice lungs after bleomycin challenge. Data are one representative experiment of two independent experiments, with 4–6 mice per group in each replicate. Graphs show mean ± SEM. Statistical significance tested by two-way ANOVA with Holm-Sidak’s multiple comparisons test; \**P* < 0.05, \*\**P* < 0.01, \*\*\**P* < 0.001. (**I**) Volcano plot of DEGs [log2 fold change (FC) > 1, adjusted *P* < 0.05] in whole lung tissue between Mr1^−/−^ and WT mice lungs at day 3 post-bleomycin challenge. The top 25 up and down-regulated genes are labelled. Horizontal line indicates *P* value threshold of 0.05. Vertical line indicates log2 fold change threshold of 1. (**J**) Top 25 significantly enriched (*P* < 0.05) pathways from Gene Ontology (GO) database (biological process) in upregulated DEGs in the lungs of Mr1^−/−^ mice compared with WT mice lungs at day 3 post-bleomycin challenge. Colour intensity indicates the statistical significance of the enrichment, and dot size signifies the number of genes upregulated in the pathway. The x axis denotes the proportion of all DEGs included in the pathway (Gene Ratio). (**K**) Heatmap depicting relative expression of selected genes of mice lungs (Day 3 and Day 0).

To elucidate the mechanism underlying the observed phenotypic differences, we evaluated immune cell infiltration between WT and Mr1^−/−^ mouse lungs using spectral flow cytometry analysis (Cytek Aurora) following a published gating strategy ^39^ (Supplementary Fig. 5B). Lung samples were collected on days 0, 3, 7 and 10 post challenge (Fig. 1D). Mr1^−/−^ mice exhibited a decreased frequency of DCs (MerTK^-^ CD11c^+^ MHII^+^) on days 3 and 7, and an increased frequency of monocytes on day 3 post bleomycin challenge. Notably, differences in the frequencies of alveolar macrophages, interstitial macrophages, neutrophils, eosinophils, NK cells, NK T cells, CD4^+^ T cells, CD8^+^ T cells, and γδ-T cells post-challenge between WT and Mr1^−/−^ mice were not statistically significant (Fig. 2, E to H; Supplementary Fig. 5, C to U).

We then investigated the transcriptomic variations between WT and Mr1^−/−^ mouse lungs. Accordingly, we obtained bulk RNA sequencing (RNA-seq) data from MAIT cell-enriched, bleomycin-challenged whole lungs. Lung samples were collected on days 0, 3, 7, 14 and 21 post challenge (Fig. 1D). Compared with WT mice, 9 (3 up, 6 down), 1729 (967 up, 762 down), 116 (54 up, 62 down), 173 (65 up, 108 down) and 192 (107 up, 85 down) genes were differentially expressed in Mr1^−/−^ mice lungs at days 0, 3, 7, 14 and 21 post-bleomycin challenge, respectively (Supplementary Fig. 5W, Fig. 2I, and data S3).

The chemokine Ccl2 (monocyte chemoattractant protein-1, MCP-1), which recruits myeloid cells towards sites of inflammation, and the proinflammatory cytokine Il6 were prominently upregulated in Mr1^−/−^ mice at day 3. Both CCL2 and IL-6 are known contributors to lung fibrosis ^40^. Gene Ontology (GO) enrichment analysis for biological processes ^41^ revealed only 3 significantly enriched (*P* < 0.05) upregulated gene sets in Mr1^−/−^ mice lungs versus WT mice lungs without bleomycin challenge (Supplementary Fig. 5X), but 1526 significantly enriched gene sets at day 3 post-bleomycin, of which the top ranked terms include leukocyte migration, regulation of cytokine production, regulation of response to external stimulus, cell adhesion, migration and chemotaxis (Fig. 2J).

Notably, expression of Xcr1, Clec9a, markers of DC ^42^ – and Flt3l and Ccr10, both essential for DC development and recruitment ^42, 43^, were significantly downregulated in Mr1^−/−^ mice relative to WT mice at day 3 post-bleomycin (Fig. 2K), consistent with downregulation of the DC population in Mr1^−/−^ mice relative to WT after bleomycin challenge shown in flow cytometry. Our data imply that MAIT cells may modulate the accumulation of immune cells in the lung after sterile lung challenge, notably DCs, during sterile lung challenges.

### Early accumulation of pulmonary CD103^+^ type 1 DCs is impaired in Mr1^−/−^ mice

We subsequently aimed to explore the subpopulations of DCs to determine which subsets contributed to the decreased accumulation observed in Mr1^−/−^ mice, following established guidelines ^44^ (Supplementary Fig. 6A). We noted an accumulation of CD45^+^ CD64^-^ CD11c^+^ MHCII^+^ cDCs (Fig. 3, A and D), particularly CD103^+^ cDC1, in lungs of WT mice on day 7 post-bleomycin. In contrast, Mr1^−/−^ mice failed to accumulate CD103^+^ DCs, with significantly lower total count and percentage of pulmonary CD103^+^ DCs at day 7 post-challenge compared to WT counterparts (Fig. 3, B and E). There was also a tendency towards impaired CD11b^+^ cDC2 accumulation in Mr1^−/−^ lungs on day 7 post-bleomycin but this did not reach statistical significance (Fig. 3, C and F).

**Fig. 3.**
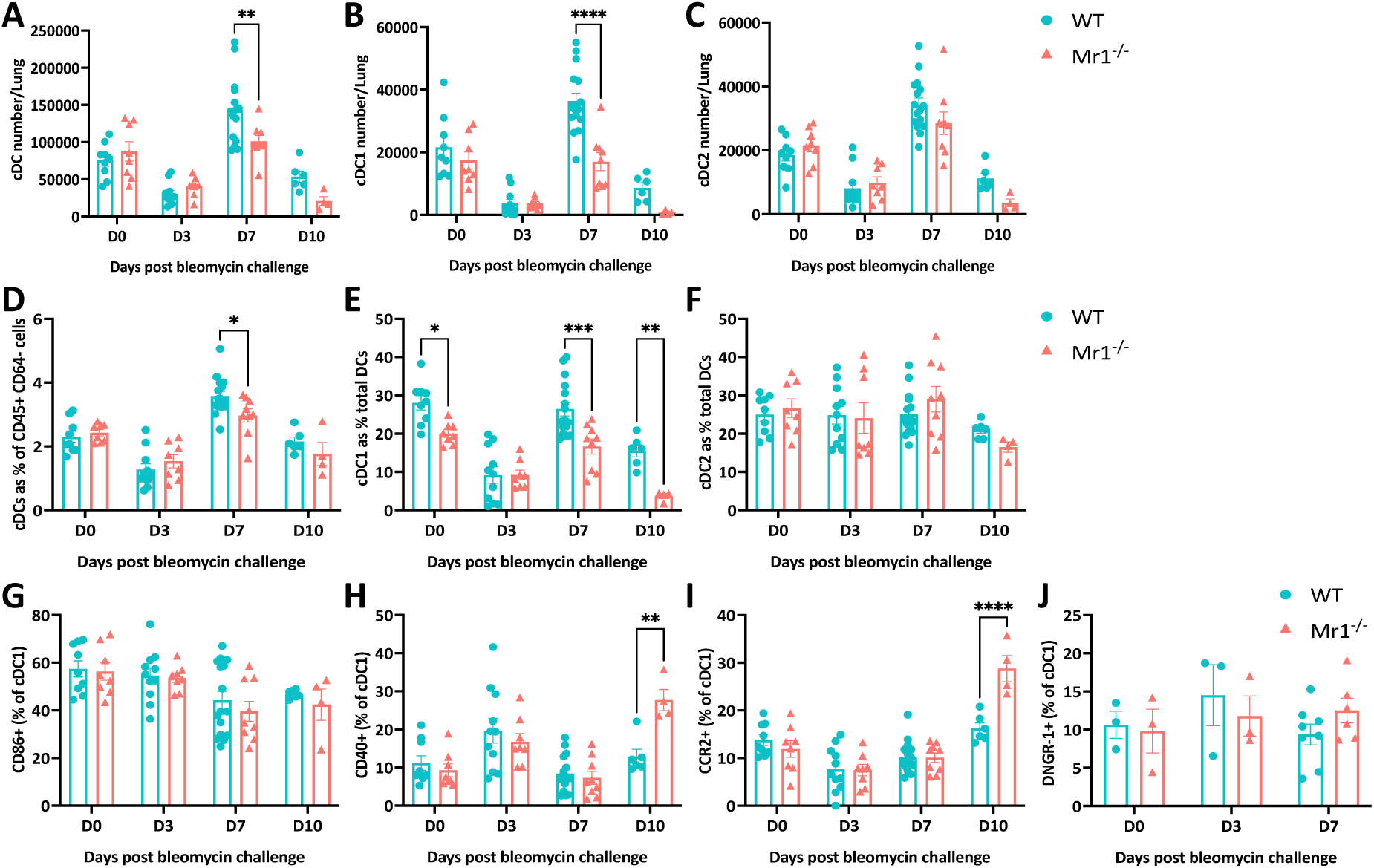
Early accumulation of CD103^+^ type 1 dendritic cells in the lungs upon bleomycin challenge is impaired in Mr1^−/−^ mice. (**A** to **C**) Absolute numbers of total cDCs (**A**), cDC1 (**B**) and cDC2 (**C**) in WT and Mr1^−/−^ mice lungs post-bleomycin challenge. (**D**) Frequencies of cDC as percentages of total CD45^+^ CD64^-^ cells in WT and Mr1^−/−^ mice lungs. (**E** and **F**) Frequencies of cDC1 (**E**) and cDC2 (**F**) as percentages of total cDC population in WT and Mr1^−/−^ mice lungs. Data represent combined data from two (Day 0, 3 and 7) or one (Day 10) independent experiments, with 3-6 mice per group. (**G** to **J**) Fraction of pulmonary cDC1 cells expressing CD86 (**G**), CD40 (**H**), CCR2 (**I**) and DNGR-1 (**J**) expressed as a percentage of pulmonary cDC1 in lungs of WT and Mr1^−/−^ mice. For CD86, CD40 and CCR2 expression on cDC1 cells, data represent combined data from two (Day 0, 3 and 7) or one (Day 10) independent experiments, with 3-6 mice per group. For DNGR-1 expression on cDC1 cells, data represent combined data from two (Day 7) or one (Day 0 and 3) independent experiments, with 2-4 mice per group. Data are presented as mean ± SEM. Significance was tested by two-way ANOVA with Sidak’s multiple comparisons test; \**P* < 0.05, \*\**P* < 0.01, \*\*\**P* < 0.001, \*\*\*\**P* < 0.0001.

To assess functional differences in cDC1 we investigated surface expression of activation markers (Supplementary Fig. 6B). No discernible difference in the expression of co-stimulatory molecules CD86 (Fig. 3G) or CD40 (Fig. 3H) was observed in the first week. Additionally, we detected no variation in the levels of CCR2, known to facilitate DC migration and recruitment ^45^, or DNGR1, a C-type lectin receptor exclusively expressed in cDC1, which is encoded by Clec9a ^46^, on cDC1 between WT and Mr1^−/−^ mice lungs at day 7 post-bleomycin (Fig. 3, I and J). However, on day 10 post bleomycin-induced lung damage, we did see an upregulation of CD40 and CCR2 in lung cDC1 cells from Mr1^−/−^ mice, suggesting that the cDC1s are showing a delayed inflammatory response in the Mr1^−/−^ mice.

### Single-cell RNA Sequencing reveals diminished cDC1 proportions in Mr1^−/−^ mice

To comprehensively delineate the cellular dynamics of major cell lineages post-bleomycin injury, we utilised single-cell RNA sequencing (scRNA-seq) (10x Genomics), Cellular Indexing of Transcriptomes and Epitopes by Sequencing (CITE-seq) and TCR sequencing. We again used the previously described MAIT-cell enriched mouse model, and obtained single cell suspensions from both WT and Mr1^−/−^ whole lungs of PBS-treated controls, as well as at days 3 and 7 post-injury, with three replicates for each time point (Fig. 4A). We obtained transcriptomes from 117,908 cells following quality control filtering. Principal component analysis highlighted variability influenced by both timepoints and mouse genotype (Supplementary Fig. 7A and B). Post-data integration and unsupervised clustering analysis, 27 cell type identities were annotated using canonical marker genes and existing scRNA-seq datasets from mouse lungs ^47–49^ (Fig. 4B, Supplementary Fig. 7C). All lineages were observed across both WT and Mr1^−/−^ mice at all three timepoints (Supplementary Fig. 7, D and E). MAIT cells exhibited a single-cell transcriptional profile akin to γδ-T cells, leading to a shared cluster in the Uniform Manifold Approximation and Projection (UMAP) (Fig. 4B). Expectedly, MAIT cells were absent in Mr1^−/−^ mice (Supplementary Fig. 7F), but there was a non-significant tendency towards an increase in CD4^+^ and CD8^+^ T cells in Mr1^−/−^ mice compared to WT mice (Supplementary Fig. 8), possibly due to a compensatory mechanism.

**Fig. 4.**
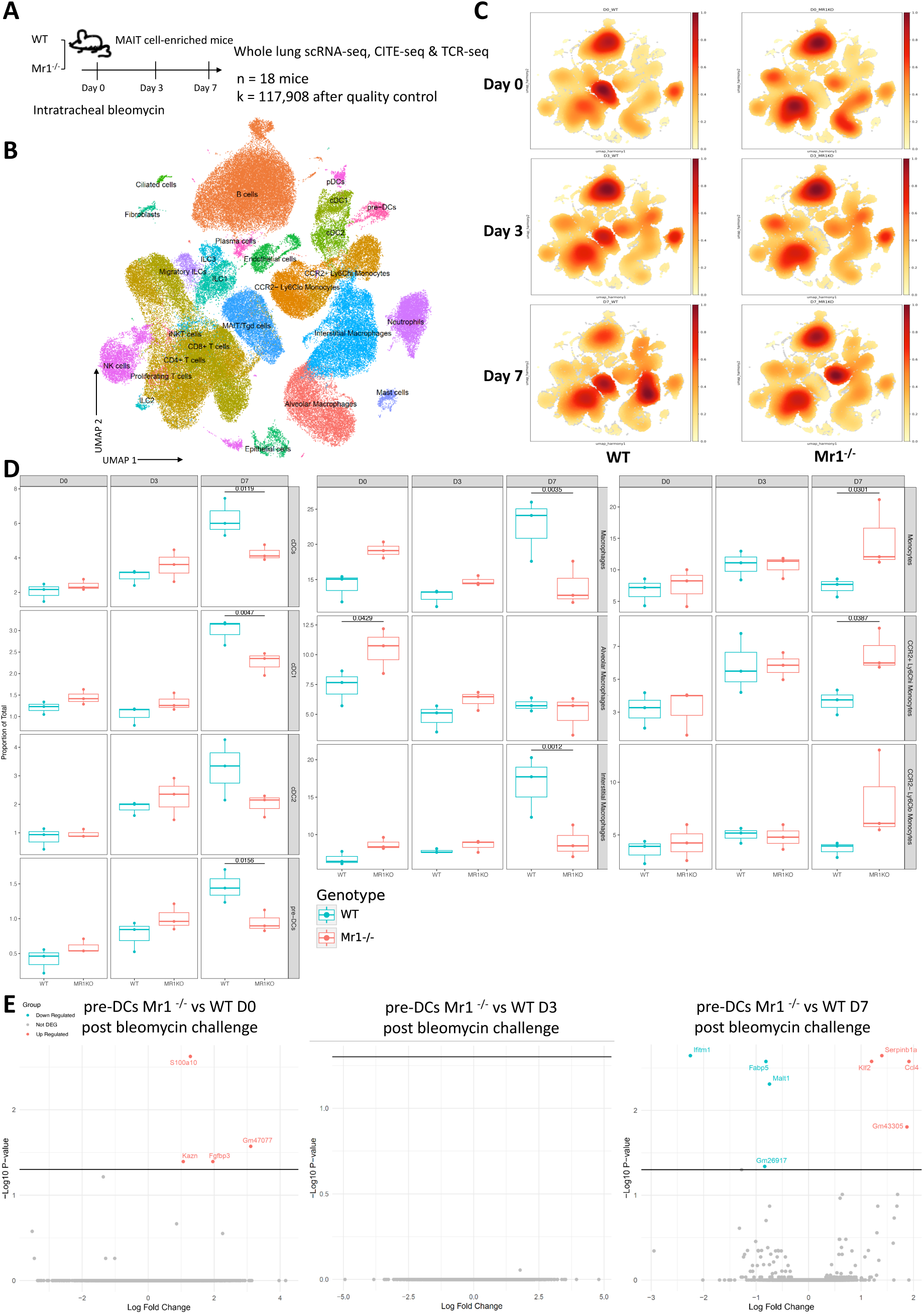
Single-cell RNA Sequencing reveals diminished cDC1 proportions in Mr1^−/−^ mice. (**A**) Single cell suspensions from whole-mouse lungs were analysed using scRNA-seq at the indicated time points after bleomycin-mediated lung injury. (**B**) UMAP embedding of 117,908 high-quality single cells colour-coded by predicted cell lineage. (**C**) UMAP embedding of cell density displaying the proportion of each cell lineage at baseline (PBS control, Day 0) and post bleomycin challenge (Day 3 and 7), stratified by timepoint and mice genotype. (**D**) Proportion of the indicated cell types of total lung cells was calculated for individual mice at the indicated time points at baseline (PBS control, Day 0) and after bleomycin challenge (Day 3 and 7) (n = 3 for each genotype). *P* values generated using a two-way ANOVA with Sidak’s multiple comparisons test. **(E**) Volcano plots showing pre-DC-specific DEGs based on adjusted *P* value < 0.05 between Mr1^−/−^ and WT mice before and post-bleomycin challenge. Horizontal line indicates *P* value threshold of 0.05.

We observed an accumulation of cDC1 (*P*=0.0047) and cDC2 (non-significant) in the lungs of WT mice but not in Mr1^−/−^ mice. These observations align with the flow cytometry data depicting a deficiency in cDC1 accumulation in Mr1^−/−^ mice (Fig. 3). Concomitantly, accumulation of monocyte and NK cells was prominently detected in the lungs of Mr1^−/−^ mice, whereas this was not the case in WT counterparts (Fig. 4, C and D; Supplementary Fig. 8). Furthermore, in the lungs of WT mice, there was a discernible expansion of both interstitial macrophages (Fig. 4D) and fibroblasts (Supplementary Fig. 8), and such expansions were absent in the Mr1^−/−^ mice.

Next we investigated the differences in cell type-specific DEGs between Mr1^−/−^ and WT mice (Supplementary Fig. 9 and data S4). Gene expression differences across most cell types were limited, with very few DEGs identified – fewer than 64 in any given cell type. Amongst pulmonary DCs and monocytes, we observed downregulation of genes in Mr1^−/−^ mice on both day 3 and 7 post-bleomycin, including Pbx1, Fcgr2b and Sh2d1b1 (Supplementary Fig. 9, data S4). The inhibitory Fc receptor Fcgr2b was consistently downregulated in Mr1^−/−^ cDC1 across all time points. This is paired with downregulation of Cxcl1/Cxcl2 in cDC2 on day 7 post-challenge (Supplementary Fig. 9), suggesting a disrupted chemokine expression profile within the cDC2 of Mr1^−/−^ mice.

We then looked at the precursor cells leading to cDCs – pre-DC population, in the lungs of both mice strains. Derived from CDP in the bone marrow, pre-DCs traffic to various tissues where they differentiate into cDC1 or cDC2, contingent upon tissue-specific and local environmental cues ^18–20^. Employing Monocle2 for trajectory analysis ^50^, we discerned a clear differentiation pathway commencing from pre-DCs (CD45^+^ MHCII^+^ CD11c^−^ Flt3^hi^ SIRP-α^−^) and culminating in cDC1 and cDC2 subsets (Supplementary Fig. 10A). During the differentiation from pre-DCs into cDC1 and cDC2, we noticed that Irf8 levels go up in cDC1 but go down in cDC2. Conversely, Irf4 levels rise in cDC2 and fall in cDC1 (Supplementary Fig. 10B). Of particular interest, Mr1^−/−^ mice exhibited a diminished pre-DC population (Fig. 4D). However, when contrasting the transcriptomic profile of pre-DCs from WT and Mr1^−/−^ mice, differential gene expression was minimal (Fig. 4E). This suggests that, during the bleomycin challenge, MAIT cells predominantly modulate the accumulation dynamics of the pre-DC population, without substantially altering their functional profile.

To test enrichment of pathways we performed GSEA analysis for all cell types across timepoints (Supplementary Fig. 11). A proinflammatory response was upregulated across various cell types in Mr1^−/−^ mice compared with WT following bleomycin. Notably, NK cells, ciliated cells, and endothelial cells exhibited this exaggerated response on day 3, while monocytes and interstitial macrophages demonstrated a similar response on day 7, which would be expected to contribute to enhanced systemic inflammation. In summary, MAIT cells predominantly influence accumulation of immune cells in response to sterile lung injury, particularly by increasing the number of cDC1s, pre-DCs and interstitial macrophages, but have more limited influence on the transcriptome of most cell types.

### Adoptive transfer of Flt3 ligand-generated bone marrow-derived dendritic cells to Mr1^−/−^ mice reverts weight loss and reduces tissue damage

We reported compromised accumulation of cDC1 in Mr1^−/−^ mice in the absence of MAIT cells. Given the exclusive presence of DNGR-1, a C-type lectin receptor, in cDC1 and its crucial role in managing tissue damage by detecting actin filaments exposed on necrotic cell death ^51^, we further probed its specific influence in our context. Previous studies have shown that DNGR-1 in DCs limits tissue damage in pancreatitis by dampening neutrophil recruitment, and DNGR-1 also controls neutrophil recruitment and pathology associated with systemic candidiasis ^52^. We therefore examined the role of cDC1-DNGR-1 in weight loss and tissue fibrosis in both WT and Mr1^−/−^ mice following bleomycin challenge. Consistent with previous data (Fig. 2, A to D), Mr1^−/−^ mice showed greater weight loss (Fig. 5A), more pronounced tissue fibrosis (Fig. 5B and Supplementary Fig. 12A), and elevated gene expression of Col1α1 and Col3α1 on D21 post-bleomycin challenge, when in comparison with WT mice (Fig. 5C and D). Significantly, weight loss and tissue fibrosis in Mr1^−/−^ mice was alleviated by intranasal adoptive transfer of Flt3L-generated bone marrow-derived dendritic cells (FLT3L-BMDC) (Supplementary Fig. 12, B to D) on day 1 post-bleomycin challenge (Fig. 5). This alleviating effect was abrogated by antibody blockade of DNGR-1, but persisted in Mr1^−/−^ mice treated with isotype control (Fig. 5). These results suggest that the protective effects offered by MAIT cells are mediated, at least partially, by regulating cDC1, and that cDC1s curb tissue damage through DNGR-1 signalling.

**Fig. 5.**
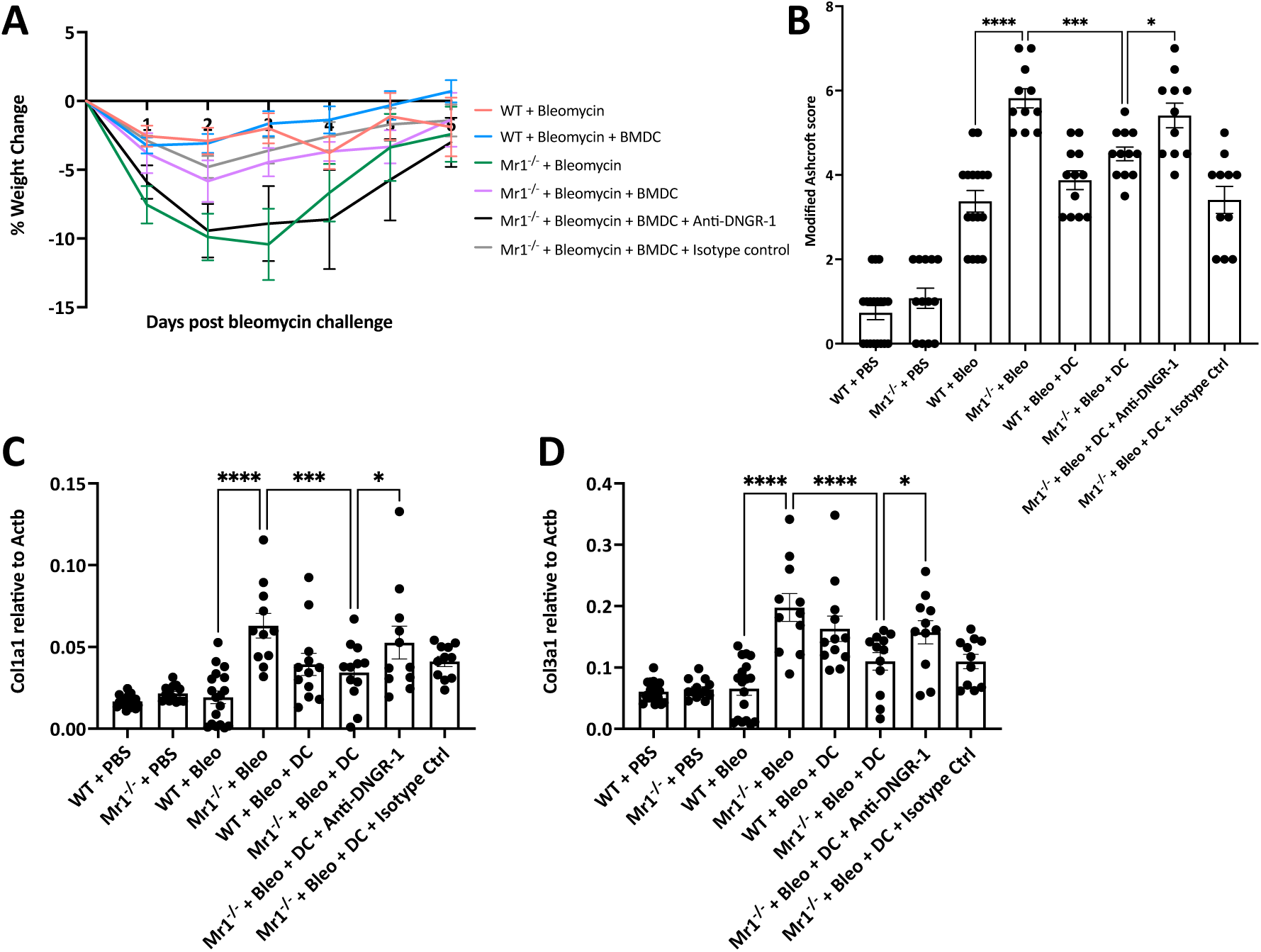
Adoptive transfer of bone marrow-derived dendritic cells (BMDC) to Mr1^−/−^ mice reverts weight loss and reduces tissue damage. (**A**) Body weight loss expressed as a percentage of the weight before bleomycin challenge. Adoptive transfer, performed twice, used 5 × 10^5^ BMDC from *Salmonella typhimurium* BRD509-infected mice bone marrows. Cells were transferred 1-day post-bleomycin challenge. (**B**) Modified Ashcroft score of lung slices of PBS or bleomycin-challenged WT and Mr1^−/−^ mice at day 21, stained with Masson’s trichrome. (**C** and **D**) Gene expression of Col1α1 (**C**), and Col3α1 (**D**) in lung homogenates of PBS or bleomycin-challenged WT and Mr1^−/−^ mice at day 21. Actb was used as a housekeeping gene. For weight loss results, data are one representative experiment of two independent experiments, with 4–6 mice per group in each replicate. For histology score and RT-qPCR results, data were pooled from two independent experiments (n=4-6 per group). Graphs show mean ± SEM. Statistical significance tested by one-way ANOVA with Holm-Sidak’s multiple comparisons test; \**P* < 0.05, \*\**P* < 0.01, \*\*\**P* < 0.001, \*\*\*\**P* < 0.0001.

### Human scRNA-sequencing datasets demonstrate differences in MAIT cell and cDC population in IPF versus non-fibrotic control lungs tissues

To compare our murine findings with human clinical data from pulmonary fibrosis, we examined published scRNA-seq datasets of IPF in the Human Lung Cell Atlas (HLCA) ^53^ and the IPF Cell Atlas^54^. MAIT cells were prominently identified only in one dataset ^55^, where 176 MAIT cells were identified in non-fibrotic controls and 141 in IPF patients (GEO reference: GSE135893) (Fig. 6A). Two datasets ^55, 56^ provided adequate participant numbers to compare cell proportions between IPF and controls (GEO reference: GSE135893 and GSE136831).

**Fig. 6.**
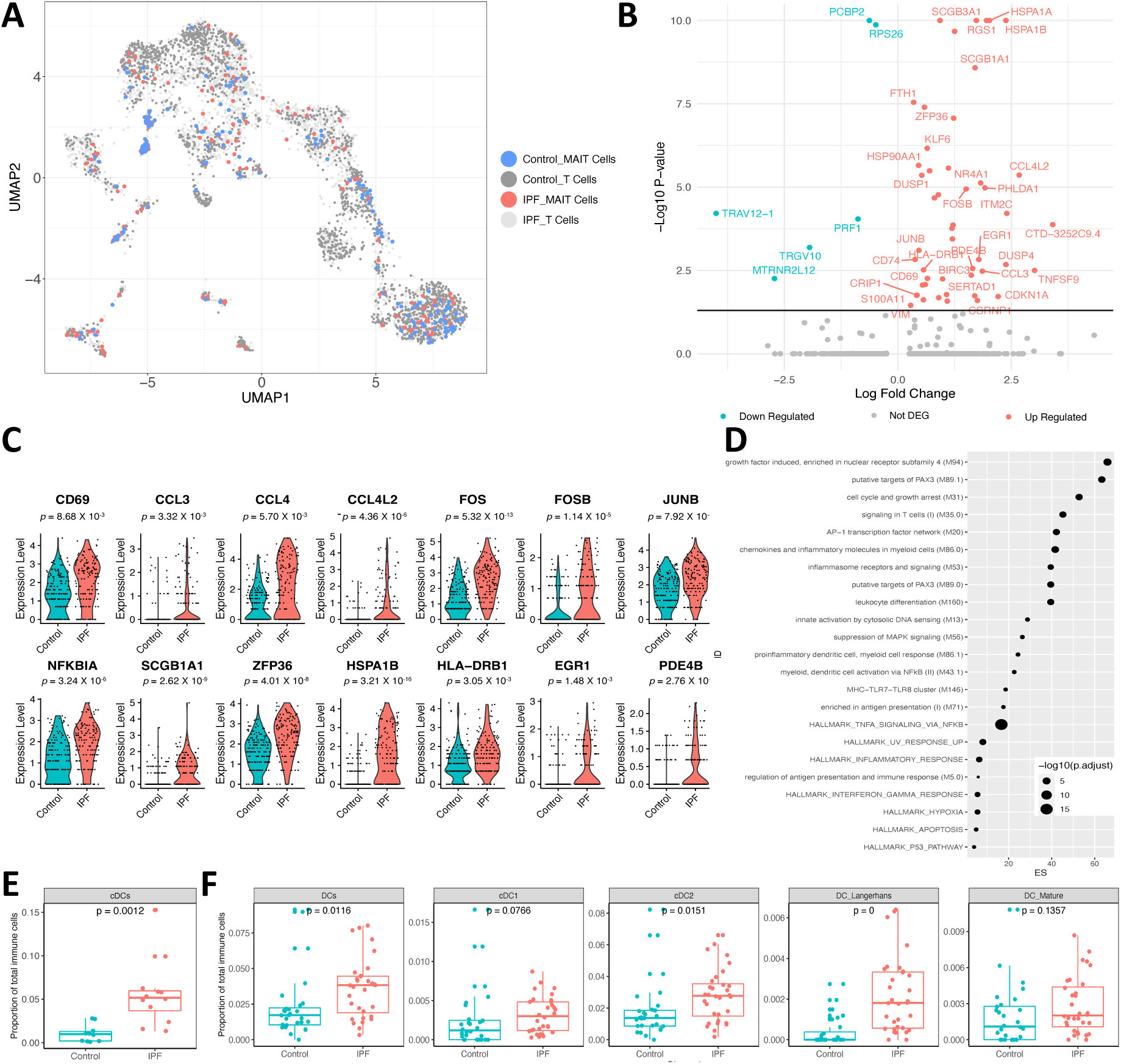
Differential gene expression and cellular frequencies in IPF patient lung MAIT cells. (**A**) UMAP showing the MAIT cells and other T cells after reintegration. (**B**) Volcano plot displays DEGs (adjusted *P* < 0.05) in IPF lung MAIT cells relative to controls. The 25 most upregulated and downregulated genes are annotated. (**C**) Violin plots illustrate expression levels of specified genes in MAIT cells, contrasting IPF with controls (GSE135893). (**D**) Analysis of DEGs for overrepresentation of blood transcriptional modules (BTM). The top 25 pathways significantly enriched (*P* < 0.05) from BTM are shown, contrasting IPF lung MAIT cells with controls. Dot size corresponds to the adjusted *P* value for each pathway, while the x-axis depicts the enrichment score (ES), calculated as the ratio of gene ratio to background ratio. (**E**) Boxplots showing the proportion of cDC relative to total immune cells in lungs of IPF patients versus controls using data from GSE135893. (**F**) Boxplots present frequencies of various DC subsets, including cDC1, cDC2, Langerhans, and mature DC, as proportions of total immune cells in lungs of IPF patients and controls, sourced from GSE136831.

MAIT cells from IPF patients’ lungs (n=10) exhibited 55 DEGs (49 up, 6 down, Supplementary Table 1) compared to controls (n=6). Notably, the MAIT cell activation marker, CD69, was among the prominently upregulated genes (Fig. 6B and C). We also observed a significant upregulation of chemokines in MAIT cells, including CCL3, CCL4, and CCL4L2, which play a pivotal role in the recruitment and activation of immune cells. Additionally, MAIT cells demonstrated enhanced expression of FOS, FOSB, and JUNB – integral components of the AP-1 transcription factor complex, which might indicate alterations in cellular signaling and responses to lung injury. The elevated expression of NFKBIA, an inhibitor of the NF-kB transcription factor, suggests potential modulations in inflammatory response pathways. And the elevated anti-inflammatory genes SCGB1A1^57^ and ZFP36^58^ in MAIT cells suggesting their modulatory role and a potential protective mechanism in the lungs of IPF patients.

We conducted an overrepresentation analysis of these DEGs using blood transcriptional modules (BTMs) ^59^. This analysis pinpointed a pronounced activation of the AP-1 transcription factor network, chemokines and inflammatory molecules in myeloid cells, as well as heightened activity related to pro-inflammatory dendritic cells and myeloid cell responses in IPF (Fig. 6D).

In the GSE135893 dataset, cDC (marked by the genes: FCER1A, CD1C, and CLEC9A) emerged as the sole cell population with a significant increase in IPF patients’ lungs (Fig. 6E and Supplementary Fig. 13B). Based on gene expression, these cDCs express CD1C, PKIB, and CLEC10A, suggesting that they are phenotypically cDC2 (Supplementary Fig. 13A). Analysis of the GSE136831 dataset revealed an elevation in the DC population within IPF patients’ lungs, with significant accumulations specifically in cDC2 (marked by FCGR2B, CLEC10A, FOXN3, ABHD12) and Langerhans cells (indicated by CD1A, FCER1A, CD1E, HLA-DQB2, S100B). There was a non-significant tendency towards an increase in cDC1 (marked by CADM1, SIPA1L3, CLEC9A, WDFY4, HDAC9) or mature DC populations (marked by CCL19, LAD1, CCR7, LAMP3, NCCRP1) (Fig. 6F and Supplementary Fig. 13C). This observation aligns with findings from our mouse lung scRNA-seq dataset, wherein DC accumulation was noted in WT mouse lungs post-bleomycin challenge, suggesting a potential role of DCs in modulating IPF-associated inflammation and fibrosis.

## DISCUSSION

In this study of sterile lung injury, we uncover a novel role for MAIT cells; they are activated and promote tissue repair, enhancing pulmonary accumulation of CD103^+^ cDC1, which limit pathology via DNGR-1. Consistent with these findings, scRNA-seq data from IPF patients reveal activated MAIT cells, and increased cDC populations in the lungs of IPF patients compared with controls. Our observations demonstrate the potential of MAIT cells as important orchestrators of tissue protection and modulators of inflammatory disease pathology.

In the context of inflammatory stimuli, MAIT cell activation is known to be triggered in at least two distinct ways: either through an MR1-TCR-dependent pathway, typically associated with antibacterial host defence, or via an MR1-TCR-independent route mediated by interleukins (IL-12/-15/-18) and type I interferon, which is often linked to antiviral responses ^34^. We observed upregulation of CD69 in lung MAIT cells of mice subjected to bleomycin challenge. This activation manifested earlier and more intensely than in non-MAIT αβ T cells, underscoring the rapid response of MAIT cells to sterile injury. There was substantial impairment in MAIT cell accumulation in both IL-18R^−/−^ and IFNαR^−/−^ mice, whilst MAIT cell CD69 upregulation was significantly impaired only in the IFNαR^−/−^ mice, consistent with our previous studies in murine influenza infection ^13^. Our findings suggest that bleomycin-induced responses are strongly cytokine-driven, with IFN-α emerging as the dominant activating cytokine in this model of sterile challenge.

After bleomycin exposure, Mr1^−/−^ mice not only suffer more severe weight loss, but also display an upregulated gene expression of Col1a1 and Col3a1, suggesting intensified tissue damage in the lungs of Mr1^−/−^ mice. Owing to their anatomical placement and resemblance to other tissue-resident lymphocytes ^36^, MAIT cells have been proposed to contribute to tissue protection ^6, 37, 60^. For instance, in the nonobese diabetic (NOD) mouse model of type 1 diabetes, Mr1^−/−^ NOD mice have displayed impaired intestinal barrier integrity, accelerated disease progression, enhanced intestinal permeability, and augmented bacterial translocation, implying a protective role of MAIT cells in maintaining surface homeostasis ^61^. Similarly, in scenarios involving allogeneic bone marrow transplantation, Mr1^−/−^ mice showed compromised survival, escalated colonic graft versus host disease (GVHD), and diminished intestinal barrier integrity^62^. These observations underscore the role of MAIT cells in preserving barrier integrity. Similar results have been observed in a mouse model of non-alcoholic steatohepatitis, where Mr1^−/−^ mice on a diet deficient in methionine and choline exhibited more severe liver damage and disease intensity compared with WT. The underlying mechanism may involve an altered ratio of pro- and anti-inflammatory hepatic macrophages ^63^, though further research is needed to determine the exact role of MAIT cells in modulating macrophage function. Moreover, a recent study showed that MAIT cells were present in the meninges and expressed high levels of antioxidant molecules, and their absence led to meningeal reactive oxygen species (ROS) accumulation and barrier leakage, highlighting the function of MAIT cells in preserving meningeal homeostasis and cognitive function ^64^. Collectively, these findings highlight the importance of MAIT cells in maintaining tissue homeostasis and reducing damage, especially in the context of inflammatory diseases.

Although MAIT cells have never been assessed in pulmonary fibrosis, murine skin resident MAIT cells exhibit a distinct tissue repair transcriptional signature^6^, similar to H2-M3 restricted CD8^+^ T cells ^35^, and seen in TCR-activated MAIT cells in humans and mice ^36, 65, 66^, indicating their local repair programme similar to other tissue resident cell types. MAIT cells have also been implicated in models of collagen-induced arthritis^67^ and chronic liver injury^68^, where their absence diminished inflammation and pathology, and their presence intensified these conditions. Pharmacological inhibition of MAIT cells alleviated fibrosis, hinting at interactions with monocytes/macrophages ^69^.

While most transcriptomic analyses in existing literature have focused on the tissue repair programme of MAIT cells under an MR1-TCR dependent activation pathway, recent published research suggests that cytokine involvement might also play a role ^7^. Our research demonstrates for the first time that both a sterile chemical insult, namely a bleomycin challenge, and a viral infection can trigger such a tissue repair programme in a cytokine-dependent manner, suggesting a functional versatility and broader applicability of the underlying mechanisms governing MAIT cell activation than previously recognised. Thus, future research would be valuable into the cytokine-dependent activation pathway and their connection to the tissue repair programme of MAIT cells, as well as the spatial relationships between MAIT cells, inflammatory and stromal cells.

Significant differences in weight loss were observed between Mr1^−/−^ and WT mice in the initial days following bleomycin challenge, yet the notable decrease in cDC1 accumulation in Mr1^−/−^ mice only occurred on day 7 post-challenge. This could indicate additional roles for MAIT cells in maintaining tissue homeostasis, in addition to the DNGR1-cDC1 mechanism elucidated here. As mentioned, our scRNA-seq dataset highlights baseline differences between Mr1^−/−^ and WT mice, notably a reduced frequency of alveolar macrophages and diminished Pbx1 expression within these cells in Mr1^−/−^ mice (Supplementary Fig. 8 and data S4). Pbx1, a key transcriptional regulator in macrophages, initiates IL10 transcription during apoptotic cell clearance via the apoptotic cell-response element (ACRE) in the IL10 promoter, indicating its significance in mitigating inflammation and maintaining tissue homeostasis during apoptosis ^70–72^. Our data suggests that MAIT cells might facilitate an anti-inflammatory state in alveolar macrophages, potentially crucial in shaping the lung’s response to tissue injury. Additionally, our total lung RNA-seq dataset highlights an early upregulation (as soon as day 3 post-bleomycin challenge) of pro-inflammatory genes, Il6 and Ccl2, in Mr1^−/−^ mice. This rapid upregulation might contribute to the early onset of weight loss observed in these mice. Further scrutiny of the scRNA-seq data identifies mast cells, interstitial macrophages, fibroblasts and CCR2^+^ Ly6C^hi^ monocytes as the primary sources of these pro-inflammatory genes (Supplementary Fig. 14).

MAIT cells and DCs coordinate in orchestrating immune responses against various pathogens and maintaining immune protection. During *Francisella tularensis* infection in mice, MAIT cell-dependent GM-CSF production contributed to monocyte differentiation into DCs, although MAIT cells were not explicitly identified as the GM-CSF source ^11^. *In vitro* investigations with human MAIT cells revealed their capacity to induce DC maturation. When co-cultured with immature human DCs in the presence of 5-Amino-6-D ribitylaminouracil/methylglyoxal (5-A-RU/MeG), MR1-dependent upregulation of CD86, CD80, CD40, and PD-L1 was observed on the DCs. Additionally, DC IL-12 production was contingent upon both MR1 and CD40L^73^. Furthermore, pulmonary MAIT cell stimulation *in vivo* with 5-A-RU/MeG or CpG led to the accumulation of CD11b^+^ DCs in the lung and migration of both CD11b^+^ and CD103^+^ DCs from the lung to the mediastinal lymph node. However this study did not explore the underlying mechanism driving the accumulation of DCs in response to MAIT cell activation^12^. We have also shown MAIT cells amplify the early immune response to adenovirus vector vaccines, a mechanism necessitating pDC-derived IFN-α, monocyte-derived IL-18, and IFN-α-induced tumour necrosis factor (TNF) from monocytes^14^. Additionally, intranasal immunisation with MAIT cell agonists and proteins activates DCs via CD40L, priming T follicular helper cells and inducing protective humoral immunity, establishing MAIT cells as potential cellular adjuvants in mucosal vaccines ^74^.

Currently, there are no *in vivo* studies examining the impact of MAIT cell and DC interactions within the context of sterile injury and tissue damage. In MAIT cell-deficient Mr1^−/−^ mice, we observed an impaired early accumulation of CD103^+^ cDC1 and pre-DC populations in the lungs upon bleomycin challenge. Interestingly, there were minimal DEGs specific to pre-DC and cDC1 when comparing WT and Mr1^−/−^ mice. This finding suggests that while MAIT cells are essential for the accumulation of cDC1, they may not modify the functional characteristics of these cells to influence their accumulation. The mechanism behind this remains to be elucidated: it is uncertain whether MAIT cells directly recruit DCs from the peripheral blood and bone marrow, indirectly draw in the precursor pre-DC population, or engage an alternative pathway. Our findings highlight that MAIT cells produce GM-CSF, IFN-γ and IL17A in the context of bleomycin-induced lung injury. Both GM-CSF and IFN-γ are important for the maturation, differentiation, and recruitment of DCs to sites of inflammation ^75, 76^. Moreover, IL-17 has been observed to augment the activation, migration, and overall functionality of airway DCs, which leads to more effective antigen-specific T cell activation and the progression of experimental allergic asthma ^77^. Our study also demonstrates an upregulation in the gene expression of Ccrl2, Ccl3, and Cxcl2 in MAIT cells following bleomycin exposure (Fig. 1G). This upregulation could be instrumental in the accumulation of pre-DC and cDC1. Notably, Ccrl2, known as an inflammatory chemokine receptor, is implicated in the trafficking of lung DCs and in managing excessive airway inflammatory responses ^78^. Ccl3, on the other hand, has been identified as a significant mobilizer of DCs, facilitating their movement into the bloodstream and toward inflammation sites ^79, 80^. We also noted a pronounced reduction in the ILC3 population in Mr1^−/−^ mice relative to WT counterparts during sterile injury, suggesting a putative role of MAIT cells in maintaining ILC3 numbers. ILC3s are known to regulate the activity of various immune cells including DCs, macrophages, eosinophils, and neutrophils, contributing to their recruitment, movement, and tissue reparative functions. In steady-state conditions, ILC3s secrete Th17-associated cytokines such as IL-17A, IL-17F, IL-22, and GM-CSF, while during inflammatory responses, they are also capable of producing IFN-γ ^81, 82^. These cytokines interact with respective receptors – IL-17R, IFN-γR, and GM-CSFR – on myeloid cells, which are particularly receptive to GM-CSF ^83, 84^. The GM-CSF produced by ILC3s has been implicated in bridging innate and adaptive immunity through its influence on myeloid cells ^85^. Thus, MAIT cells, through the production of cytokines such as GM-CSF, IFN-γ and IL17A, along with various chemokines, may exert a direct influence on the migration and functionality of DCs. Additionally, these factors could indirectly affect the function and maintenance of ILC3s, subsequently affecting the downstream accumulation and activity of diverse myeloid populations in response to sterile lung injury.

Our research provides evidence that cDC1 cells contribute to tissue protection by mediating responses to cellular damage through DNGR-1 signaling. DNGR-1, or CLEC9A, a type II transmembrane protein categorized within the C-type lectin receptor (CLR) family’s group V, is abundantly expressed on cDC1 cells. This receptor has a unique role in recognizing dead cells by binding with its C-type lectin-like domain (CTLD) to F-actin, which becomes exposed upon disruption of cell membrane integrity ^51^. The subsequent phosphorylation of a hemITAM, especially at the tyrosine 7 residue, leads to SYK activation ^86–88^. This triggers an intracellular response involving activation of NADPH oxidase and production of ROS, which induces lipid peroxidation, damages phagosomal membranes and causes the release of their contents into the cytosol ^87, 89^. Consequently, this permits cross-presentation of antigens to CD8^+^ T cells through the MHC-I pathway, traditionally used for presenting endogenous antigens. The ability of DNGR-1-mediated cDC1 signaling to detect and respond to cell damage not only mitigates the risk of excessive inflammation and collateral damage but also facilitates the prompt resolution of the immune response, underlying the critical role of cDC1s in adaptive immunity and tissue homeostasis.

Our study indicates that MAIT cells are notably activated in the lungs of IPF patients compared to non-fibrotic controls. To date, there has been limited literature examining the function of MAIT cells in IPF. One notable study reported decreased counts of MAIT cells in the peripheral blood of IPF patients, suggesting a potential migration to the lungs ^90^. Future research could delve into the involvement of MAIT cells in the onset and progression of pulmonary fibrosis. Another promising area of exploration is the interaction between MAIT cells and lung microbiota during fibrosis, given the known responsiveness of MAIT cells to bacterial and viral infections. Evaluating MAIT cells as potential therapeutic targets through the use of synthetic blocking or activating ligands may also yield valuable insights.

Our study has several limitations. Firstly, while we identified the central role of MAIT cells in the accumulation of CD103^+^ cDC1 and pre-DC populations, the specific mechanisms through which MAIT cells influence this accumulation remain ambiguous. The exact interplay between MAIT cell-produced cytokines, such as GM-CSF and IFN-γ, and the accumulation of DC requires further elucidation. The potential interactions and crosstalk between MAIT cells and other myeloid cells also warrant a deeper investigation to comprehensively understand their combined roles in shaping immune responses to sterile challenge. Secondly, in contrast to the early accumulation of cDC1 in the lungs of WT mice subjected to bleomycin challenge, human IPF patient data shows a more pronounced accumulation of cDC2, indicating species differences and the need for careful interpretation in the context of the chronic nature of human IPF and the dynamics of DC infiltration and function. Thirdly, the intricate processes of tissue damage and fibrosis, which involve numerous structural cells such as epithelial cells and fibroblasts, present difficulties for us to study. The methods we used, like flow cytometry or scRNA-seq, inherently lose information about structural cells and spatial resolution during single cell preparation, leaving various questions unanswered: How are MAIT cells spatially distributed and interacting within the lung tissue? Do their gene expression pattern and intercellular signalling pathways vary across regions, correlating with tissue damage? These questions could be answered through spatial transcriptomic approaches.

Employing the bleomycin-challenged murine model, a well-accepted surrogate of interstitial lung disease (ILD) pathogenesis ^91^, we have identified a crucial role for MAIT cells in counteracting lung injury. This offers invaluable insights into the complex mechanisms underpinning pulmonary tissue repair. Specifically, airway damage and tissue remodelling emerge as common pathological traits across a wide spectrum of respiratory disorders, significantly contributing to their progression. While acting as a marker for disease progression, tissue remodelling is also a contributor to severe clinical symptoms and disease severity, causing impaired quality of life^92^. These effects of remodelling are prominent in conditions such as ILD, pneumonia, and acute respiratory distress syndrome^93^. Given the pathological importance of remodelling in lung diseases our findings are potentially important for the development of novel antifibrotic therapeutic strategies. For instance, the protective activities of MAIT cells could be enhanced by use of MAIT cell stimulating ligands, either synthetically produced, or administered through riboflavin-competent commensal organisms^94^, or through targeting the DNGR-1 pathway. Future studies will be required to further delineate the precise roles of MAIT cells in the context of human lung injury and repair, and to validate the translational potential of our findings in real-world clinical settings, but in summary we have for the first time generated *in vivo* data of MAIT cells protecting against lung damage, independently of a bacterial or viral pathogen.

## MATERIALS AND METHODS

### Mice model and *in vivo* bleomycin challenge

C57BL/6 mice (aged 8–10 weeks) were purchased from University of Oxford Biomedical Services (BMS), Charles River or Envigo. Mr1^−/−^ mice ^95^ (kindly provided by Dr Claire Hutchings, University of Oxford, MGI ID: 3664578), Il18r1^tm1Aki^ mice (kindly provided by Prof Kevin Maloy, University of Oxford, MGI: 2136765), and Ifnar1^tm1Agt^ mice (kindly provided by Dr Claire Hutchings, University of Oxford, MGI ID: 1930950) were bred in house and used at 8-10 weeks of age. Sex and age were matched between groups. All mice were housed in specific pathogen-free conditions. For indicated experiments, C57BL/6 and Mr1^−/−^ mice were co-housed for ≥ 28 days to normalize the microbiome between strains ^62^. All work was performed under UK Home Office license PPL P61FAD253 in accordance with the UK Animal (Scientific Procedures) Act 1986. For bleomycin challenge, mice were anaesthetized with isoflurane then treated intratracheally with 1.875 U/Kg (mice weight) of bleomycin sulphate (Apollo Scientific, Cat. No. BI3543) in 50 μL of PBS.

### Mice pulmonary MAIT cell expansion using *Salmonella Typhimurium* BRD509

*S. typhimurium* BRD509 were prepared as previously described ^13^. Mice were infected intranasally with 10^6^ CFU *S. typhimurium* BRD509 in 50 μL PBS under isoflurane anaesthesia.

### Generation of MR1 tetramers

Murine MR1-5-OP-RU and MR1-6-FP monomers were provided by the NIH Tetramer Facility. Tetramers were generated using Brilliant Violet 421 (BV421)-Streptavidin and Phycoerythrin (PE)-Streptavidin (BioLegend, Cat. No. 405225 and 405245, respectively) following the NIH Tetramer Facility’s guidelines.

### Antibodies staining for flow cytometry and cell sorting

Murine lung tissues were prepared as described previously ^36^. For measurement of intracellular markers, 1 × Brefeldin A (eBioscience™, Cat. No. 00-4506-51) was added 4 hours before staining. Lung cells were blocked with anti-Fc receptor 2.4G2 and/or 6-FP tetramer for 15 min at room temperature (RT), stained with viability dye, fluorescently-labelled MR1 tetramer and/or flow cytometric antibodies for 20 min at RT. Staining antibodies, clones and concentrations are listed in Supplementary Table 2. Samples were washed in FACS buffer (PBS + 0.5% BSA + 2 mM EDTA), and cells were fixed for 15 min RT using IC fixation buffer then washed twice with 1 × permeabilization buffer (eBioscience^TM^, Cat. No. 00-8222-49 and 00-8333-56, respectively). Intracellular staining was performed overnight at 4°C. Samples were subsequently washed twice and stored in FACS buffer at 4°C until analysed on BD LSRII flow cytometer.

For live cell sorting on murine lung MAIT cells, lung single-cell suspension was purified with a 40%: 70% Percoll gradient. The sorting was conducted on a BD Aria III directly into a 350µL lysis buffer (Buffer RLT plus, supplemented with 10µL β-ME per 1mL Buffer RLT plus from the QiaGen RNeasy Plus Micro Kit, Cat. No. 74034) and subsequently stored at -80 oC for future batch RNA extraction.

For multiparameter spectral flow cytometry analysis, we used the “AF as a tag” (AF) function in the SpectroFlo (Cytek Biosciences, CA) software ^96^. 6 unique AF tags were disassociated from unstained mouse lung samples and included in the unmixing strategy.

### Total RNA extraction and RNA integrity assessment

RNA extraction was performed by single column centrifugation using the RNeasy® Plus Micro Kit (Qiagen, Cat No. 74034) following the manufacturer’s protocol. RNA integrity was assessed by Agilent High Sensitivity RNA ScreenTape Assay on an Agilent 4200 TapeStation following the manufacturer’s protocol (Agilent, Cat. No. 5067-5579, 5580 and 5581).

### mRNA isolation, library preparation, sequencing by Novogene

Total RNA was subsequently submitted to Novogene following their sample preparation and shipping instructions. All further laboratory work was performed by Novogene using their commercial protocol. RNA was reverse transcribed and cDNA amplified by *in vitro* transcription with the TaKaRa SMART-Seq v4 Ultra Low Input RNA Kit for Sequencing (TaKaRa, Cat. No. 634889) or a low input method using NEB Next® Ultra RNA Library Prep Kit for Illumina® (NEB, Cat. No. 7530). First-strand cDNA synthesis and tailing by reverse transcription were performed using SMART (Switching Mechanism at 5’ End of RNA Template) technology. Following first-strand synthesis, cDNA was amplified by PCR to produce the library. Quality control of the library was performed by quantification with a Qubit 2.0 fluorimeter and by qPCR. Insert size was measured by the Agilent 2100 Bioanalyzer automated gel electrophoresis system. The library was sequenced using the NovaSeq platform with Illumina sequencing technology to generate 150bp paired-end reads.

### RNA-sequencing data analysis

NovaSeq platform images were first converted into raw sequence reads via Illumina’s CASAVA software, stored in FASTQ format. After filtering out low-quality and adapter reads, the remaining clean reads were mapped using STAR version 2.6.1 ^97^ against the mus musculus GRCm38 reference genome (GenBank accession number GCA_000001635.2). Successful mapping was determined by a rate over 70%, with the results preserved as BAM files ^98^. Read quantification involved featureCounts 1.5.0 ^99^, which converted BAM files to a table of gene IDs and counts per sample. Differential expression analysis was performed in in R (version 4.1.0) using DESeq2 (version 1.30.1) ^100^. DEGs were defined as log2 fold-change > 1 and adjusted *P* < 0.05. VennDiagram (version 1.6.20). And ggplot2 (version 3.2.1), pheatmap (version 1.4.3) and ggrepel (version 0.8.1) were used for data visualization. clusterProfiler (version 4.0) ^101^ and Mfuzz ^102^ are used for GO enrichment and time-series analysis, respectively. GSEA was performed using GSEA software (version 4.1.0) ^103^. Bulk RNA-seq data cell type deconvolution was performed utilising MUlti-Subject SIngle Cell deconvolution (MuSiC) method ^104^. For this analysis, cell-type-specific gene expression data from a bleomycin-induced lung injury mouse model, acquired from the single-cell RNA-seq dataset (GSE141259) published by Strunz. M. *et al.* was used as the reference dataset ^49^.

### Propagation, quantification and mice infection of PR8 influenza A virus

Propagation, quantification and mice infection of PR8 influenza A virus were performed as previously described ^33^.

### 10x Genomics library generation, sequencing and computational analysis

Sequencing libraries were generated using 10x Genomics Chromium Next GEM Single Cell 5’ Reagent kit v2 (Dual Index) following manufacturer’s instructions (CG000330 Rev D). ADT-labelled (BioLegend, Cat. No. 155861, 155863 and 199903) cells were loaded onto the Chromium iX (10x Genomics) at a concentration of ∼1 x 10^6^ cells/mL, with 50,000 cells loaded per channel. One channer was loaded per 2 mice lung samples. Library generation was performed using Biomek FX^P^ Laboratory Automation Workstation (Beckman Coulter) at MRC Weatherall Institute of Molecular Medicine Single-Cell Facility (WIMM, University of Oxford). Library quality and concentration was assessed using a Bioanalyzer (Agilent) and Qubit 2.0 Fluorometer (Thermo Fisher Scientific), respectively. Libraries were sequenced on an Illumina NovaSeq 6000 to a mean depth of 40,000 read pairs/cell for scRNA-seq, 10,000 read pairs/cell for Cite-seq and TCR-seq, performed at Novogene.

10x Genomics cellranger analysis pipelines were used to generate single cell gene counts. Reads from gene expression and TCR library were aligned to the mouse mm10 reference genome (version 2020-A) and GRCm38 Mouse V(D)J Reference-7.0.0 (May 17, 2022), respectively, and quantified using cellranger multi pipeline together with those from ADT library.

Hashtag oligo (HTO) data underwent a transformation using Seurat’s centred log-ratio (CLR) transformation ^105^. Demultiplexing of HTO hashtags was subsequently performed manually with CITEviz ^106^, followed by normalisation of CITE-seq data via the dsb package ^107^. Ambient RNA was removed using decontX ^108^. The RNA-seq data were then processed using Scanpy ^109^, which involved doublet removal with Scrublet ^110^ and cell filtering with specified thresholds for total counts (1,000 – 60,000), genes by counts (500 – 6,000), and mitochondrial counts (0 – 10%). The filtered data underwent normalization to achieve a total sum of 10,000 and were log-transformed with a pseudocount of 1. Highly variable genes were identified by setting the flavour to “cell_ranger”. Principal Component Analysis (PCA) was conducted, with the number of principal components set using the KneeLocator function. Cells from different mice were subsequently integrated using the Harmony algorithm ^111^. Neighbourhoods were identified with n_neighbors set to 5, followed by dimensionality reduction with UMAP ^112^ and partitioning cell type with Leiden clustering at a resolution of 2.0 ^113^. DEGs between cell types were identified using the rank_genes_groups function with a t-test. Cell clusters were identified using both RNA and protein expression data. T cells were further selected for reintegration and subset identification using the same method as previously stated, except that the clustering resolution was set to 1.0.

PCA on the overall transcriptome for each mouse was based on pseudobulk counts, computed by summing the counts of all cells in each mouse. Genes with low expression was filtered using filterByExpr in edgeR (version 3.36.0) ^114^ by setting min.count to 3, and using model matrix adjusted for time point and genotype. Normalisation factors were calculated using calcNormFactors in edgeR and the pseudocounts were then normalised using voom in limma (version 3.50.3) ^115^.

MAIT cells and iNKT cells were identified using clonotypes.csv and filtered_contig_annotations.csv, based on the output generated by cellranger multi pipeline. A cell is designated as a MAIT cell if it is part of a clonotype that exhibits Trav1 and Traj33 expression. A cell is designated as an iNKT cell if it is part of a clonotype that exhibits Trav11 and Traj18 expression.

Differential gene expression between Mr1^−/−^ and WT mice across various time points and cell types was analysed using DESeq2 ^100^ and pseudobulk counts, computed by summing the counts of all cells within each cell type for each mouse. Enriched gene sets were identified using the pre-ranked gene-set enrichment analysis (GSEA) algorithm implemented in the FGSEA R package ^116^. Genes were ranked with the log2 fold change for the relevant coefficient calculated by DESeq2. Enrichment was assessed with gene set list from MSigDB’s Hallmark collection.

In the trajectory analysis of DC populations, Monocle 2 ^50^ (version 2.22.0) was utilized. The raw count data was processed to establish a CellDataSet object within Monocle 2 by setting the expressionFamily to a negative binomial distribution with a fixed variance. Cell ordering was achieved using genes identified by dpFeature. Dimension reduction for visualization was carried out using DDRTree. Pre-cDCs were identified as the root_state during the cell ordering process. Trajectories were generated independently for cells from different groups. The expression profiles of selected genes in the two differentiation branches (cDC1 and cDC2) were visualized using the “plot_genes_branched_heatmap” function within the Monocle2 package ^50^.

### Analysis of human IPF scRNA-seq datasets in HLCA and IPF cell atlas

Cells within the HLCA and IPF cell atlas were assessed for TRAV1-2 expression. Those exhibiting positive TRAV1-2 expression levels were categorized as MAIT cells. The primary identification of MAIT cells was in GSE135893 ^55^, and only these MAIT cells were utilized in subsequent analyses. Differential gene expression between MAIT cells from IPF patients and those from healthy controls was determined using the FindMarkers function in Seurat ^105^, with a pseudocount set to 1. For the overrepresentation test, modules in BTM ^59^ were employed, utilizing the enricher function in clusterProfiler ^101^.

Both GSE135893 and GSE136831 ^55, 56^ were analysed in terms of immune cell composition changes, given that both datasets contained more than three samples from the IPF and healthy control groups, respectively. Marker genes for DC populations were derived from the original publication.

### Culture and adoptive transfer of Flt3 ligand-generated bone marrow-derived dendritic cells

B6.SJL-Ptprc^a^ Pepc^b^/BoyJ mice were used as donor mice and all donor mice were infected with 10^6^ CFU *S. typhimurium* BRD509 4 weeks before harvesting bone marrow cells. FLT3L-BMDC were generated by culture in RPMI complete containing murine Flt3L at 200ng/mL and murine GM-CSF at 20ng/mL as previously described ^117^. DNGR-1 was highly expressed in the MHCII^+^ CD11c^+^ CD24^hi^ subsets of Flt3L BMDCs (Supplementary Fig. 10), which correspond to the CD103^+^ subset of lung DCs. CD11c^+^ FLT3L-BMDCs were enriched using anti-PE microbeads (Miltenyi Biotec, Cat. No. 130-048-801). 5 × 10^5^ FLT3L-BMDCs were given into each recipient Mr1^−/−^ mouse intranasally ^118^ at day 1 post-bleomycin challenge. When necessary, mice were treated i.p. with 100 μg of 7H11 anti-DNGR-1 blocking antibody or isotype-matched control (BioXCell, Cat. No. BE0305 and BE0088, respectively). Injections were administered daily from day -1 to day 10 post-bleomycin challenge.

### Histology

The left lobes of the mice lungs were preserved in 10% neutral buffered formalin, sequentially dehydrated with an ethanol gradient, cleared with Histo-Clear II, and infiltrated with paraffin wax. Subsequently, paraffin-embedded sections (4 μm thick) of these lobes were stained with Masson’s trichrome (Abcam, Cat. No. ab150686) following manufacturer’s instructions. To evaluate the extent of fibrosis, the modified Ashcroft scoring system was employed for a semiquantitative analysis ^119^.

### Hydroxyproline assay

Hydroxyproline was measured using 10 mg of lung tissue using a hydroxyproline assay kit (Sigma-Aldrich, Cat. No. MAK008) per the manufacturer’s instructions.

### RNA quantification, purity check and reverse transcription

For RT-qPCR experiments, total lung RNA was extracted as described above. RNA quantity and quality were assessed using a Nanodrop 2000 (Thermo Scientific) following the manufacturer’s protocol. Isolated RNA was converted to cDNA in preparation for qPCR using a High-Capacity cDNA Reverse Transcription Kit with RNase Inhibitor (Applied Biosystems, Cat. No. 4374966) following manufacturer’s protocol. Template RNA and reagents were thawed on ice. The reverse transcription reaction mix was prepared and incubated in the Programmable Thermal Controller as the following steps: Step 1: 25°C, 10 minutes; Step 2: 37°C, 120 minutes; Step 3: 85°C, 5 minutes; Step 4: 4°C, ∞.

### Reverse transcription quantitative polymerase chain reaction (RT-qPCR)

For qPCR reactions, 2 × QuantiFast SYBR Green PCR Master Mix kit (QiaGen, Cat No. 204056) was used following the manufacturer’s instructions. PCR reaction mix was prepared, mixed, and appropriate volumes were dispensed into the wells of a PCR plate. Template cDNA was added to the individual wells containing the reaction mix. qPCR plate was in loaded into a Bio-Rad CFX96. qPCR was performed following the manufacturer (Bio-Rad)’s instructions. Thermal cycling conditions were set up as the following steps: PCR initial heat activation: 95°C, 5 minutes; 2-step cycling: Denaturation: 95°C, 10s; Combined annealing/extension: 60°C, 30s; 35-40 cycles in total. For all tests, *P* < 0.05 was considered statistically significant.

### Data analysis and statistics

Flow cytometry data were acquired on a BD LSRII Flow Cytometer (BD Biosciences) or Cytek Aurora (Cytek Biosciences) and processed in FlowJo version 10.8.1 (FlowJo, LLC). All data was analysed in Prism version 9.2.0 (GraphPad) and RStudio version 1.4.1717. For *in vivo* mouse data analysis, various tests were deployed as required, including unpaired t tests, Mann-Whitney tests, one-way ANOVA with Dunnett’s or Sidak’s multiple comparisons, Kruskal-Wallis with Dunn’s multiple comparisons, and two-way ANOVA with Sidak’s multiple comparisons, Holm-Sidak’s multiple comparisons or Fisher’s LSD test. A *P* value less than 0.05 was considered significant.

## Acknowledgements

We thank S. B. Morgan for the guidance to X.Z. during the initial stage of this work; H. Ferry for help with cell sorting; A. Byrne for support with the bleomycin mice model; the NIH Tetramer Facility for the MR1 tetramers; A. Davison from Cytek Biosciences for assistance with the analysis of spectral flow cytometry data. This work was funded by China Scholarship Council (CSC) – Nuffield Department of Medicine (NDM) award (X.Z) and grants from the Wellcome Trust (104553/z/14/z, 211050/Z/18/z to T.S.C.H. and 222426/Z/21/Z P.K.).

## Author contributions

T.S.C.H., P.K. and X.Z. jointly conceived the work. X.Z. performed all the mice experiments. X.Z., W.L. and M. G. performed the single-cell RNA sequencing experiment. X.Z., S.L., T.S.C.H. and P.K. performed the data analysis. X.Z. drafted the manuscript, and all authors contributed to editing of the manuscript.

