## Supplementary Materials for "MAIT cells protect against sterile lung injury"

Supplemental Materials File

**This file includes:**

Supplementary Figs. 1 to 14

Supplementary Tables 1 and 2

Captions for Supplementary Data 1 to 4

**Other Supplementary Materials for this manuscript include the following:**

Supplementary Data 1 to 4

Supplementary Data 1 – DEGs of MAIT cells, WT bleomycin-challenged versus WT unchallenged

Supplementary Data 2 – DEGs of MAIT cells, WT PR8 virus infected versus WT B6 uninfected

Supplementary Data 3 – DEGs of total lung, $Mr1^{-/-}$ versus WT B6

Supplementary Data 4 – DEGs, cell type-specific, WT versus $Mr1^{-/-}$

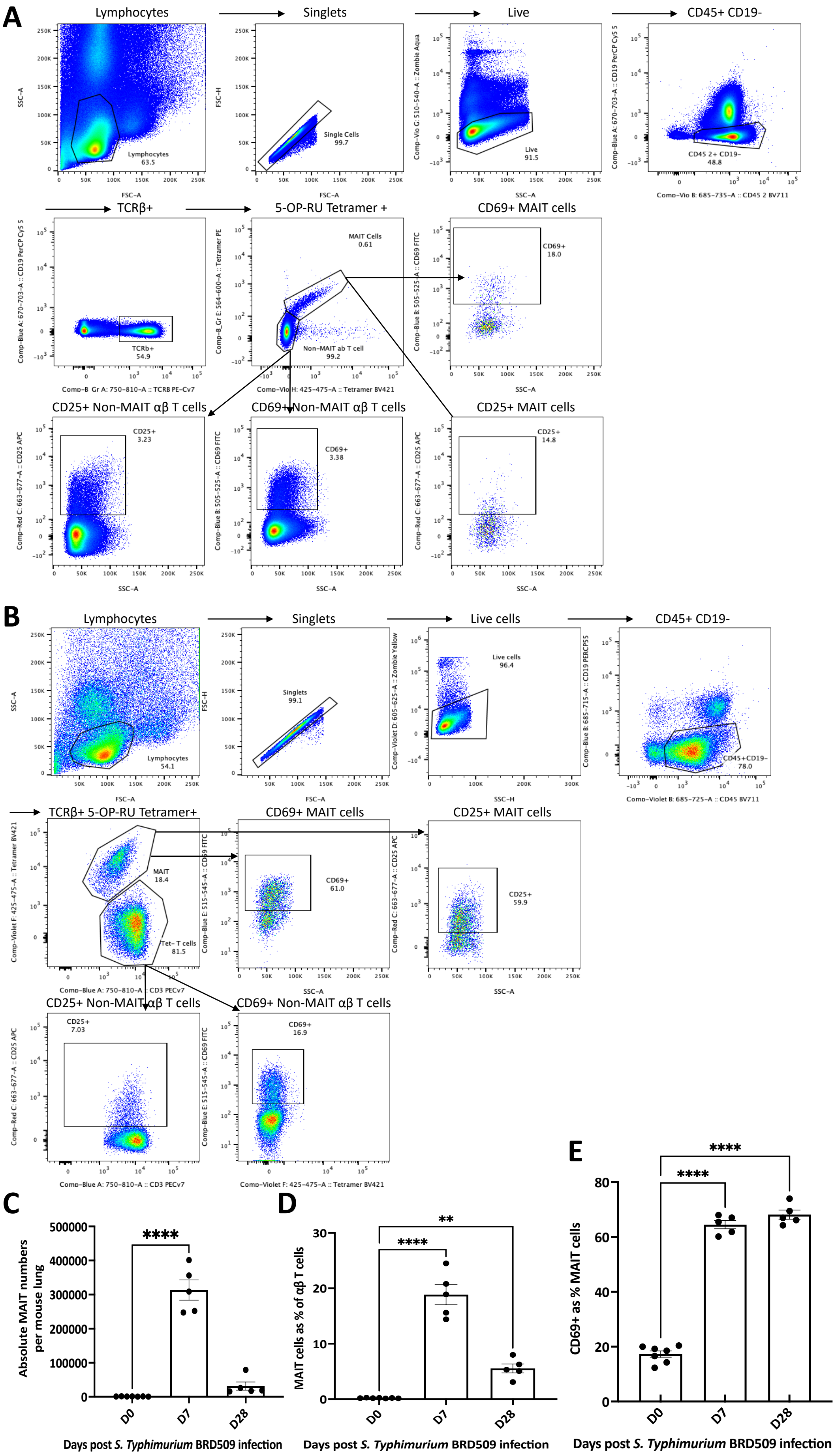
A
Lymphocytes
Singlets
Live
CD45+ CD19-
Lymphocytes 63.5
Single Cells 99.7
Live 91.5
CD45 2+ CD19- 48.8
FSC-A
SSC-A
FSC-H
TCRβ+
5-OP-RU Tetramer +
CD69+ MAIT cells
TCRb+ 54.9
MAIT Cells 0.61
Non-MAIT ab T cell 99.2
CD69+ 18.0
CD25+ Non-MAIT αβ T cells
CD69+ Non-MAIT αβ T cells
CD25+ MAIT cells
CD25+ 3.23
CD69+ 3.38
CD25+ 14.8
B
Lymphocytes
Singlets
Live cells
CD45+ CD19-
Lymphocytes 54.1
Singlets 99.1
Live cells 96.4
CD45+CD19- 78.0
TCRβ+ 5-OP-RU Tetramer+
CD69+ MAIT cells
CD25+ MAIT cells
MAIT 18.4
Tet- T cells 81.5
CD69+ 61.0
CD25+ 59.9
CD25+ Non-MAIT αβ T cells
CD69+ Non-MAIT αβ T cells
CD25+ 7.03
CD69+ 16.9
C
Absolute MAIT numbers per mouse lung
****
D0
D7
D28
Days post S. Typhimurium BRD509 infection
D
MAIT cells as % of αβ T cells
****
**
D0
D7
D28
Days post S. Typhimurium BRD509 infection
E
CD69+ as % MAIT cells
****
****
D0
D7
D28
Days post S. Typhimurium BRD509 infection

**Supplementary Fig. 1. Temporal dynamics of MAIT cells in pulmonary immune response to *S. typhimurium* infection.** (**A** and **B**) Gating scheme for the identification of MAIT cells (TCRβ$^+$ CD45.2$^+$ CD19$^-$ MR1-5-OP-RU tetramer$^+$ cells), and expression of CD25 and CD69 on MAIT and non-MAIT αβ T cells in naïve mice lung (**A**) and *S. typhimurium* BRD509 infected mice lung (**B**). (**C** and **D**) Bar plots showing the absolute MAIT cell numbers (**C**) and percentage of MAIT cells of total pulmonary αβ T cells (**D**) post BRD509 infection. (**E**) Proportion of pulmonary MAIT cells expressing CD69 expressed as a percentage of pulmonary MAIT cells post BRD509 infection. Data (mean ± S.E.M.) are one representative experiment of two independent experiments, with 5–7 mice per group in each replicate. Statistical significance tested by one-way ANOVA with Dunnett's multiple comparison tests. ** $P < 0.01$; **** $P < 0.0001$.

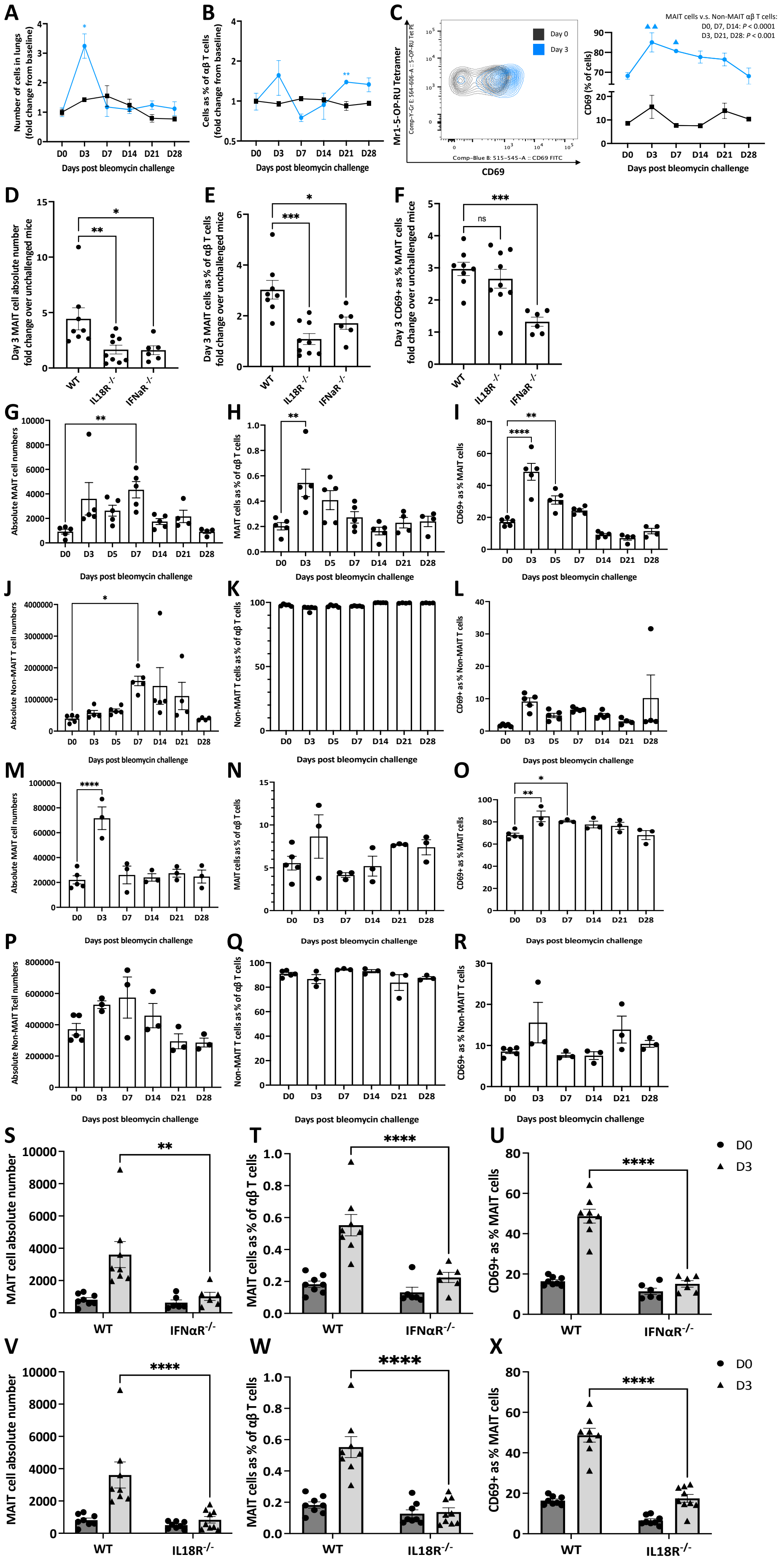

A
Number of cells in lungs (fold change from baseline)
Days post bleomycin challenge
B
Cells as % of αβ T cells (fold change from baseline)
Days post bleomycin challenge
C
Mr1-5-OP-RU Tetramer
CD69
Day 0
Day 3
MAIT cells v.s. Non-MAIT αβ T cells:
D0, D7, D14: P < 0.0001
D3, D21, D28: P < 0.001
CD69 (% of cells)
Days post bleomycin challenge
D
Day 3 MAIT cell absolute number fold change over unchallenged mice
WT
IL18R -/-
IFNaR -/-
E
Day 3 MAIT cells as % of αβ T cells fold change over unchallenged mice
F
Day 3 CD69+ as % MAIT cells fold change over unchallenged mice
ns
G
Absolute MAIT cell numbers
Days post bleomycin challenge
H
MAIT cells as % of αβ T cells
I
CD69+ as % MAIT cells
J
Absolute Non-MAIT T cell numbers
K
Non-MAIT T cells as % of αβ T cells
L
CD69+ as % Non-MAIT T cells
M
Absolute MAIT cell numbers
N
MAIT cells as % of αβ T cells
O
CD69+ as % MAIT cells
P
Absolute Non-MAIT Tcell numbers
Q
Non-MAIT T cells as % of αβ T cells
R
CD69+ as % Non-MAIT T cells
S
MAIT cell absolute number
WT
IFNαR-/-
T
MAIT cells as % of αβ T cells
U
CD69+ as % MAIT cells
D0
D3
V
MAIT cell absolute number
WT
IL18R-/-
W
MAIT cells as % of αβ T cells
X
CD69+ as % MAIT cells
D0
D3

**Supplementary Fig. 2. Cytokine-modulated accumulation and activation of pulmonary MAIT cells upon bleomycin stimulation.** Accumulation and activation of lung MAIT cells in WT C57BL/6 mice or mice with IL18R or IFNαR deficiency following a bleomycin challenge. (**A**) Fold change in the absolute MAIT cell and non-MAIT αβ T cell number in the lungs of **MAIT cell-boosted mice** (infected with *S. Typhimurium* BRD509 four weeks prior to bleomycin challenge) post-bleomycin challenge, relative to unchallenged D0 controls (PBS). Comparisons between MAIT cell subsets and non-MAIT αβ T cell subsets at individual time-points, were conducted using unpaired t tests or Mann-Whitney tests, *$P < 0.05$, **$P < 0.01$, ***$P < 0.001$, ****$P < 0.0001$. (**B**) Fold change in MAIT cell and non-MAIT αβ T cell frequency as the percentage of total pulmonary αβ T cells in the lungs of **MAIT cell-boosted mice** post-bleomycin challenge, relative to unchallenged controls. Comparisons between MAIT cell subsets and non-MAIT αβ T cell subsets at individual time-points, were conducted using unpaired t tests or Mann-Whitney tests, *$P < 0.05$, **$P < 0.01$, ***$P < 0.001$, ****$P < 0.0001$. (**C**) Proportion of pulmonary MAIT cells and non-MAIT αβ T cells expressing CD69. Statistical comparisons across different timepoints post-bleomycin challenge and unchallenged PBS control were made using one-way ANOVA with Dunnett's or Dunn's multiple comparison tests, $^{\blacktriangle}P < 0.05$, $^{\blacktriangle\blacktriangle}P < 0.01$, $^{\blacktriangle\blacktriangle\blacktriangle}P < 0.001$, $^{\blacktriangle\blacktriangle\blacktriangle\blacktriangle}P < 0.0001$. Comparisons between MAIT cell subsets and non-MAIT αβ T cell subsets at individual time-points, were conducted using unpaired t tests or Mann-Whitney tests. The data are presented as the mean ± SEM of a single experiment, with 3-5 mice in each group. (**D**) Fold change in the number of pulmonary MAIT cells in naïve WT or cytokine knock out mice on day 3 post-bleomycin challenge, relative to unchallenged controls. (**E**) Fold change in the frequency of MAIT cells as a percentage of total pulmonary αβ T cells in naïve WT or cytokine knock out mice on day 3 post challenge, relative to unchallenged controls. (**F**) Fold change in the proportion of pulmonary MAIT cells expressing CD69 in naïve WT or cytokine knock out mice on day 3 post challenge, relative to unchallenged controls. For Supplementary Fig. 2 D to F, differences in significance between WT mice and cytokine deficient mice were assessed using one-way ANOVA with Dunnett's multiple comparison tests. Data are mean ± SEM, pooled from three independent experiments, with 2–5 mice per group in each replicate; *$P < 0.05$, **$P < 0.01$, ***$P < 0.001$, ****$P < 0.0001$. (**G** to **L**) Raw absolute number data for **Fig.1**, **A to C**. Statistical comparisons across different timepoints post-bleomycin challenge and unchallenged PBS control were made using one-way ANOVA with Dunnett's or Dunn's multiple comparison tests, *$P < 0.05$, **$P < 0.01$, ***$P < 0.001$, ****$P < 0.0001$. (**M** to **R**) Raw absolute number data for **Supplementary Fig. 1, A to C**. Statistical comparisons across different timepoints post-bleomycin challenge and unchallenged PBS control were made using one-way ANOVA with Dunnett's or Dunn's multiple comparison tests, *$P < 0.05$, **$P < 0.01$, ***$P < 0.001$, ****$P < 0.0001$. (**S** to **X**) Raw absolute number data for **Supplementary Fig. 1, D to F**. Statistical significance tested by two-way ANOVA with Holm-Sidak's multiple comparisons test; *$P < 0.05$, **$P < 0.01$, ***$P < 0.001$, ****$P < 0.0001$.

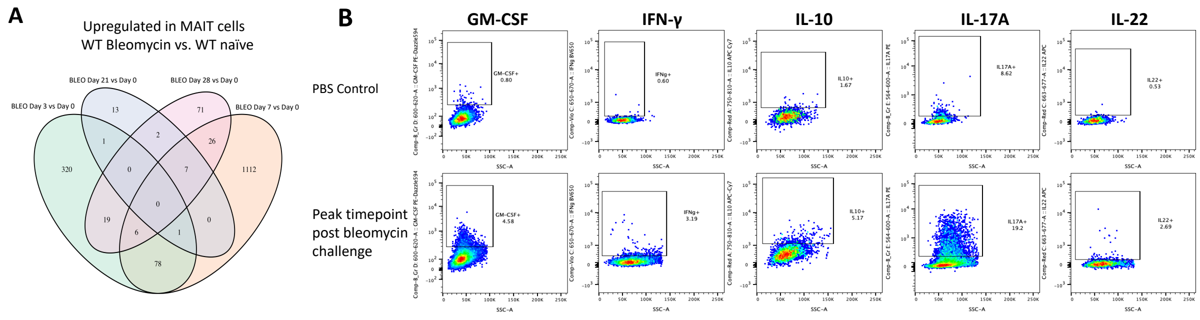


**Supplementary Fig. 3: Gene expression and cytokine profile of pulmonary MAIT cells following bleomycin-induced lung injury.** (**A**) Venn diagram showing the overlap of genes up-regulated in mice lung MAIT cells on day 3, 7, 21 and 28 post-bleomycin challenge, compared to unchallenged controls. (**B**) Flow cytometry plot illustrating the expression levels of GM-CSF, IFN-γ, IL-10, IL-17A, and IL-22 in lung MAIT cells from MAIT-cell enriched mice after bleomycin challenge. Peak response times to bleomycin for each cytokine are presented, with Day 10 for GM-CSF, IFN-γ, and IL-10, and Day 7 for IL-17A and IL-22.

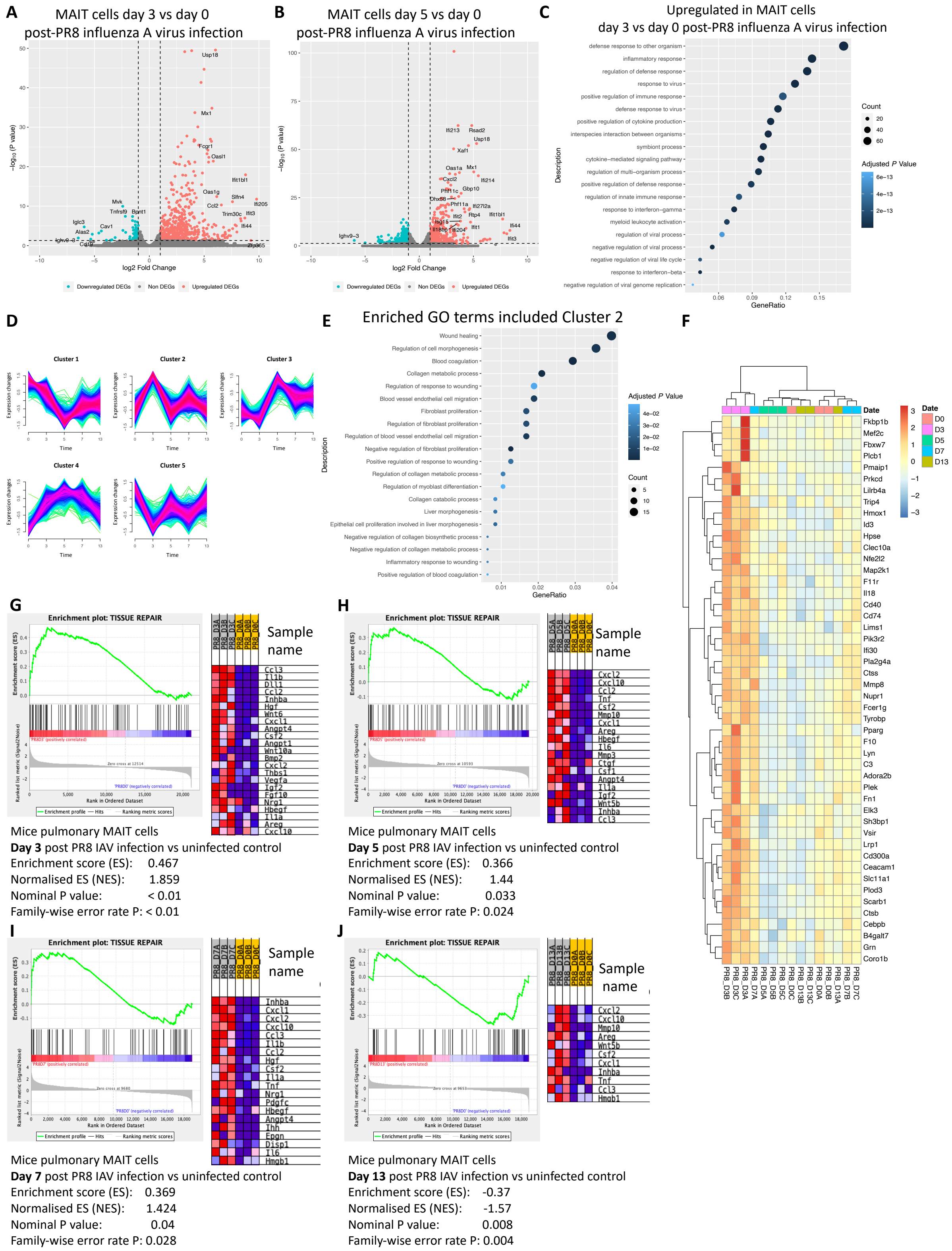

**Supplementary Fig. 4. PR8 influenza A virus (IAV) infection activates tissue repair programme of lung MAIT cells**. (**A** and **B**) Volcano plot of differentially expressed genes (DEGs) [log2 fold change (FC) > 1, adjusted *P* < 0.05] in mice lung MAIT cells on day 3 (**A**)

and 5 (**B**) post-PR8 IAV infection compared to uninfected controls. The top 10 up and down-regulated genes are annotated. Horizontal line indicates *P* value threshold of 0.05. Vertical line indicates log2 fold change threshold of 1. (**C**) Top 25 significantly enriched ($P < 0.01$) pathways from Gene Ontology (GO) database (biological process) in upregulated DEGs of mice lung MAIT cells at day 3 post-PR8 IAV infection compared to uninfected controls. (**D** to **F**) Time-series analysis showed the gene alteration trends of PR8 IAV-infected mice MAIT cells at five different time points. (**D**) Time-series expression of five gene clusters obtained from the time-series analysis. The different colours indicated the degree of match between changes of gene expression and the major changes of the clusters. Red, blue and green represented high, moderate and low match degrees, respectively. (**E**) Tissue repair-related GO terms (biological process) enriched from genes in cluster 2. (**F**) Heatmap depicting relative expression of genes associated with tissue repair at different time points. (**G** to **J**) Gene set enrichment analysis (GSEA) for tissue repair gene signature [1] in mouse MAIT cells. Heatmap showing expression of leading-edge subset genes within the gene set (red, highest expression; blue, lowest) for mouse pulmonary MAIT cells at day 3 (**G**), 5 (**H**), 7 (**I**) and 13 (**J**) post-PR8 IAV infection and uninfected control. For dot plot (**C** and **E**), the colour intensity indicates the statistical significance of the upregulation, with the dot size representing the number of genes upregulated in each pathway. The x-axis illustrates the proportion of all differentially expressed genes included in each pathway (Gene Ratio).

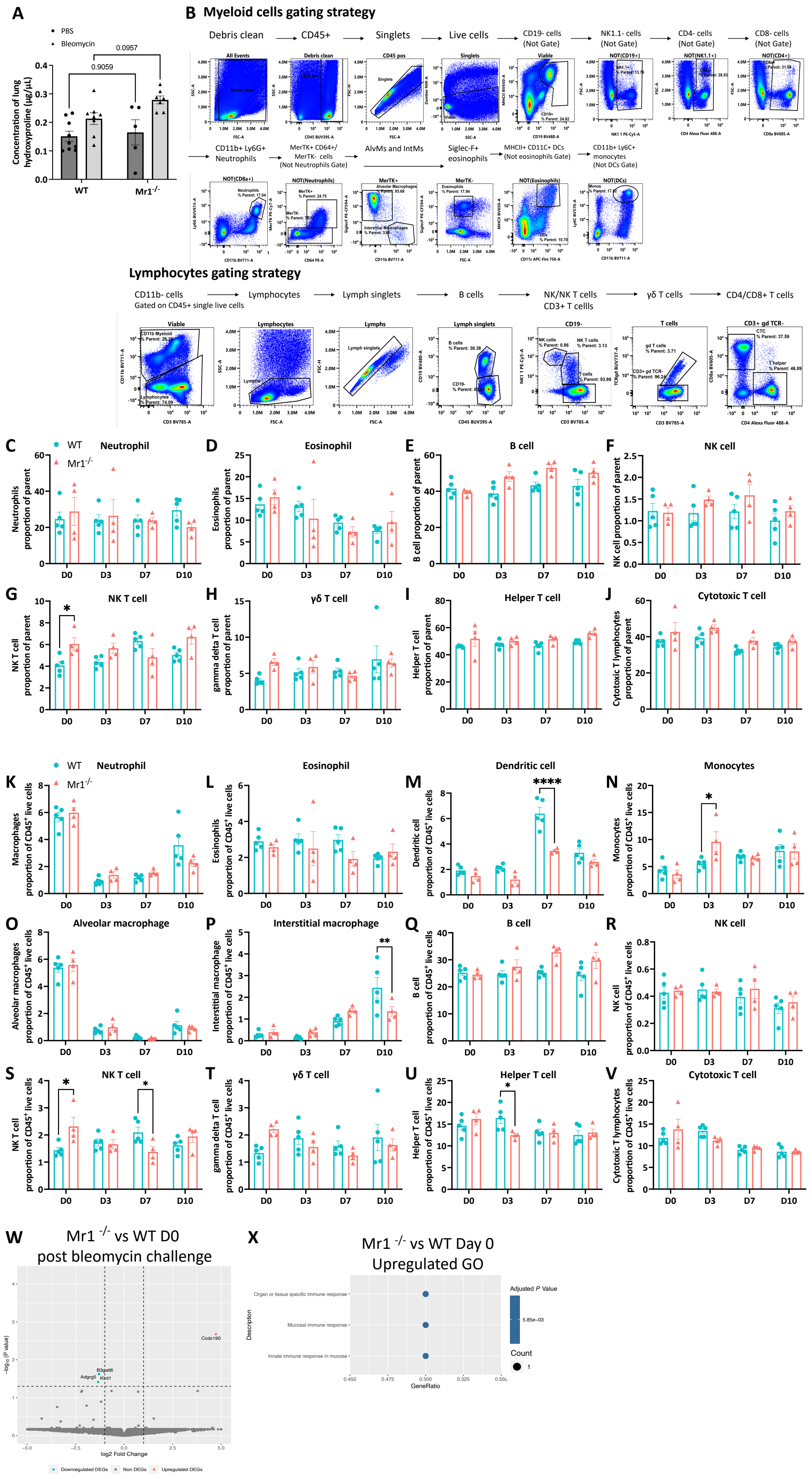
A
PBS
Bleomycin
0.0957
0.9059
Concentration of lung hydroxyproline (μg/μL)
WT
Mr1-/-
B Myeloid cells gating strategy
Debris clean
CD45+
Singlets
Live cells
CD19- cells (Not Gate)
NK1.1- cells (Not Gate)
CD4- cells (Not Gate)
CD8- cells (Not Gate)
CD11b+ Ly6G+ Neutrophils
MerTK+ CD64+/ MerTK- cells (Not Neutrophils Gate)
AlvMs and IntMs
Siglec-F+ eosinophils
MHCII+ CD11C+ DCs (Not eosinophils Gate)
CD11b+ Ly6C+ monocytes (Not DCs Gate)
Lymphocytes gating strategy
CD11b- cells Gated on CD45+ single live cells
Lymphocytes
Lymph singlets
B cells
NK/NK T cells CD3+ T cellls
γδ T cells
CD4/CD8+ T cells
C Neutrophil
WT
Mr1-/-
Neutrophils proportion of parent
D Eosinophil
Eosinophils proportion of parent
E B cell
B cell proportion of parent
F NK cell
NK cell proportion of parent
G NK T cell
NK T cell proportion of parent
H γδ T cell
gamma delta T cell proportion of parent
I Helper T cell
Helper T cell proportion of parent
J Cytotoxic T cell
Cytotoxic T lymphocytes proportion of parent
K Neutrophil
Macrophages proportion of CD45+ live cells
L Eosinophil
Eosinophils proportion of CD45+ live cells
M Dendritic cell
Dendritic cell proportion of CD45+ live cells
N Monocytes
Monocytes proportion of CD45+ live cells
O Alveolar macrophage
Alveolar macrophages proportion of CD45+ live cells
P Interstitial macrophage
Interstitial macrophage proportion of CD45+ live cells
Q B cell
B cell proportion of CD45+ live cells
R NK cell
NK cell proportion of CD45+ live cells
S NK T cell
NK T cell proportion of CD45+ live cells
T γδ T cell
gamma delta T cell proportion of CD45+ live cells
U Helper T cell
Helper T cell proportion of CD45+ live cells
V Cytotoxic T cell
Cytotoxic T lymphocytes proportion of CD45+ live cells
D0
D3
D7
D10
W Mr1 -/- vs WT D0 post bleomycin challenge
Ccdc190
B3galt6
Adgrg5
Klrd1
-log10 (P value)
log2 Fold Change
Downregulated DEGs
Non DEGs
Upregulated DEGs
X Mr1 -/- vs WT Day 0 Upregulated GO
Organ or tissue specific immune response
Mucosal immune response
Innate immune response in mucosa
Description
GeneRatio
Adjusted P Value
5.85e-03
Count

**Supplementary Fig. 5. Pulmonary cellular dynamics and gene expression in both WT and Mr1$^{-/-}$ mice following bleomycin challenge.** (**A**) Hydroxyproline concentration in WT and Mr1$^{-/-}$ mice at D21 after bleomycin. Data are one representative experiment of two independent experiments, with 5–9 mice per group in each replicate. Statistical significance tested by two-way ANOVA with Holm-Sidak's multiple comparisons test. (**B**) Flow cytometry gating strategy for the identification of pulmonary myeloid cells and lymphocytes. Representative flow cytometry plots showing gating for cells isolated from mouse lung tissue to identify neutrophil (CD11b$^+$ Ly6G$^+$), eosinophil (MerTK$^-$ Siglec-F$^+$), alveolar macrophage (MerTK$^+$ CD64$^+$ CD11b$^-$ Siglec-F$^+$), interstitial macrophage (MerTK$^+$ CD64$^+$ CD11b$^+$ SiglecF$^-$), dendritic cell (MerTK$^-$ CD11c$^+$ MHII$^+$), monocyte (MerTK$^-$ CD64$^+$ CD11b$^+$ Ly6C$^+$), B cell (CD3- CD19+), CD4$^+$ T cell (CD3$^+$ CD19$^-$ CD4$^+$), CD8$^+$ T cell (CD3$^+$ CD19$^-$ CD8$^+$), NK cells (CD3$^-$ NK1.1$^+$), NK T cell (CD3$^+$ NK1.1$^+$), and γδ-T cell (TCRγδ$^+$) populations. (**C** to **J**) Frequencies of neutrophil (**C**), eosinophil (**D**), B cell (**E**), NK cell (**F**), NK T cell (**G**), γδ-T cell (**H**), helper T cell (**I**) and cytotoxic T cell (**J**) **as percentages of parent** in WT and Mr1$^{-/-}$ mice lungs. (**K** to **V**) Frequencies of neutrophil (**K**), eosinophil (**L**), dendritic cell (**M**), monocyte (**N**), alveolar macrophage (**O**), interstitial macrophage (**P**), B cell (**Q**), NK cell (**R**), NK T cell (**S**), γδ-T cell (**T**), helper T cell (**U**) and cytotoxic T cell (**V**) **as percentages of total live immune cells** (Live CD45$^+$ cells) in WT and Mr1$^{-/-}$ mice lungs. Data are one representative experiment of two independent experiments, with 4–6 mice per group in each replicate. Graphs show mean ± SEM. Statistical significance tested by two-way ANOVA with Holm-Sidak's multiple comparisons test; *$P < 0.05$. (**W**) Volcano plot of DEGs [log2 fold change (FC) > 1, adjusted $P < 0.05$] in whole lung tissue from WT and Mr1$^{-/-}$ mice lungs without bleomycin challenge (Day 0). The top 25 up and down-regulated genes are labelled. Horizontal line indicates $P$ value threshold of 0.05. Vertical line indicates log2 fold change threshold of 1. (**X**) All significantly enriched ($P < 0.05$) upregulated pathways from Gene

Ontology (GO) database (biological process) in upregulated DEGs in the lungs of Mr1$^{-/-}$ mice compared with WT mice lungs without bleomycin challenge (Day 0). Colour intensity indicates the statistical significance of the upregulation, and dot size signifies the number of genes upregulated in the pathway. The x axis denotes the proportion of all DEGs included in the pathway (Gene Ratio).

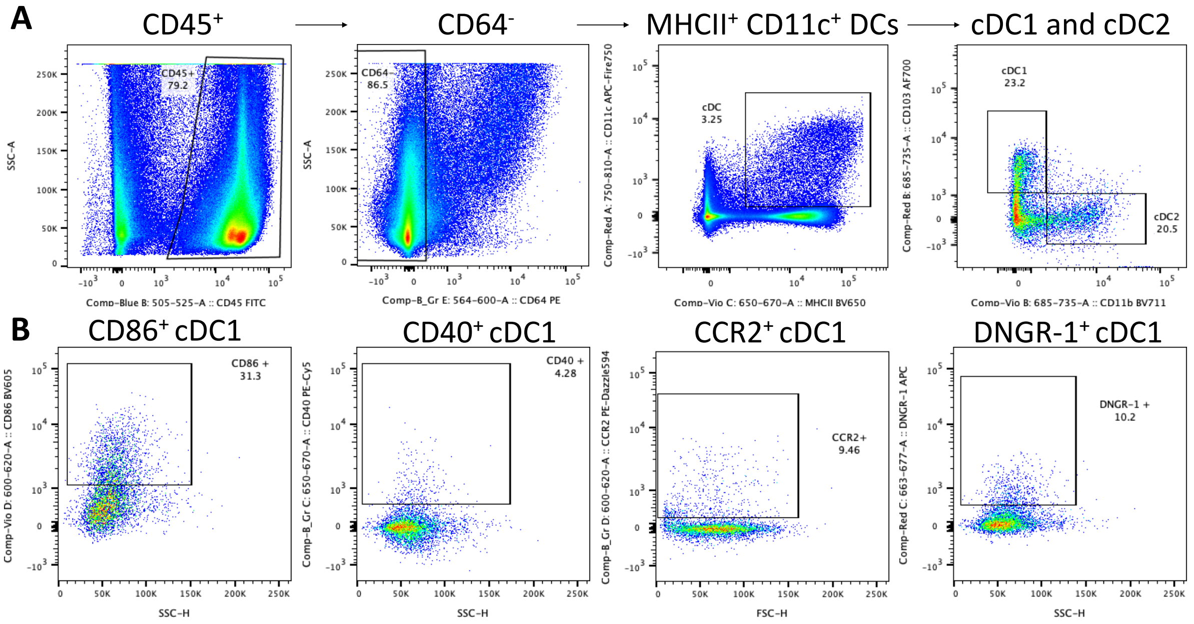


**Supplementary Fig. 6. Characterisation and activation status of mouse lung conventional dendritic cells subsets.** (**A**) Representative flow cytometry plots of lung conventional dendritic cells (cDC), type 1 cDCs (cDC1) and type 2 cDCs (cDC2). (**B**) Representative flow cytometry plots showing expression of CD86, CD40, CCR2 and DNGR-1, gated on live pulmonary CD45$^{+}$ CD64$^{-}$ CD11c$^{+}$ MHCII$^{+}$ CD103$^{+}$ cDC1 cells.

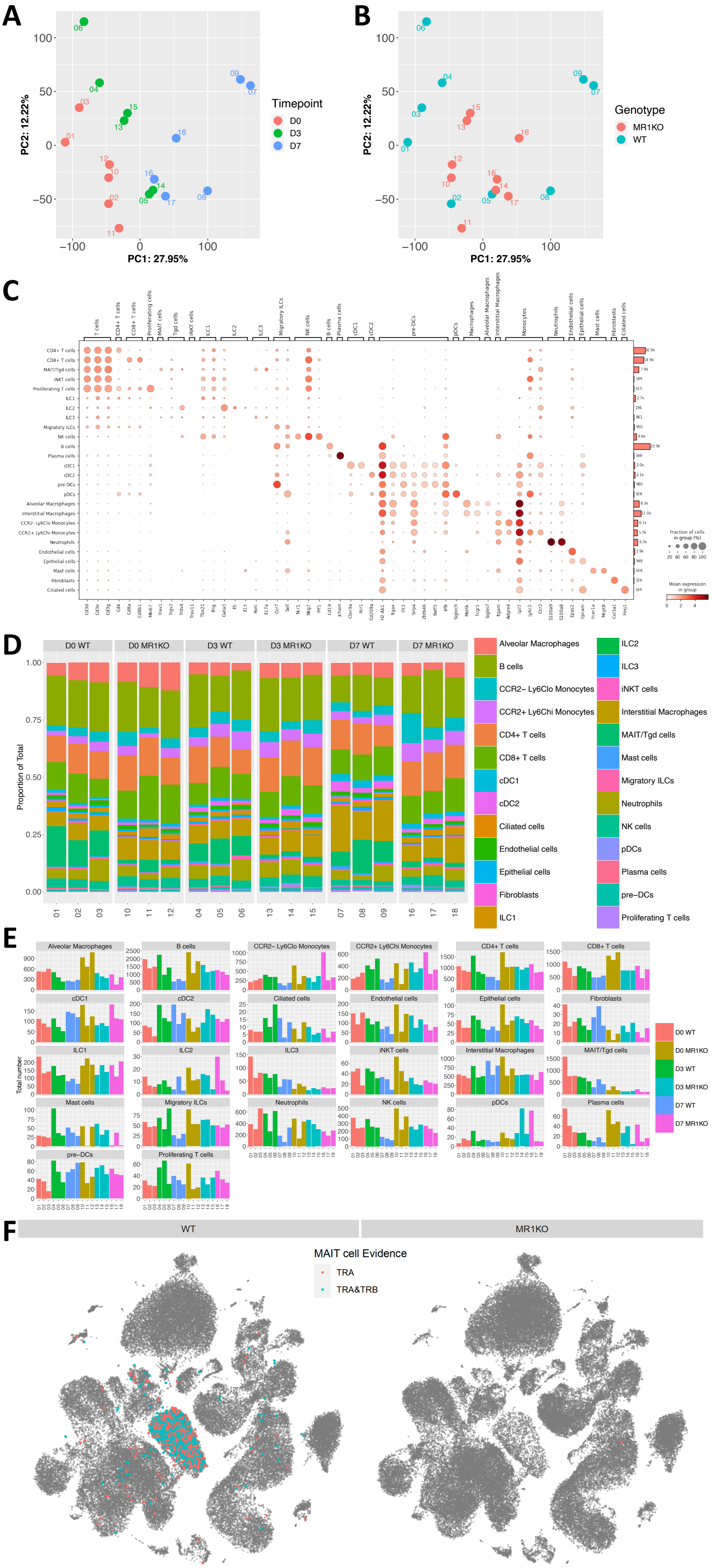

A
B
PC1: 27.95%
PC2: 12.22%
Timepoint
D0
D3
D7
Genotype
MR1KO
WT
C
D
D0 WT
D0 MR1KO
D3 WT
D3 MR1KO
D7 WT
D7 MR1KO
Proportion of Total
Alveolar Macrophages
B cells
CCR2− Ly6Clo Monocytes
CCR2+ Ly6Chi Monocytes
CD4+ T cells
CD8+ T cells
cDC1
cDC2
Ciliated cells
Endothelial cells
Epithelial cells
Fibroblasts
ILC1
ILC2
ILC3
iNKT cells
Interstitial Macrophages
MAIT/Tgd cells
Mast cells
Migratory ILCs
Neutrophils
NK cells
pDCs
Plasma cells
pre−DCs
Proliferating T cells
E
Total number
F
WT
MR1KO
MAIT cell Evidence
TRA
TRA&TRB

**Supplementary Fig. 7. Differential gene expression and cellular distribution in mouse lung cell lineages across genotypes and timepoints post-challenge.** (**A** and **B**) Principal component analysis on the overall transcriptomes of each sample coloured by timepoint (**A**) or mice genotype (**B**). (**C**) Dot plot showing the top discriminative gene sets for each cell lineage compared with every other lineage. Dot size represents fraction of cells in each group. Colour scales denote the normalised mean gene expression for each cluster. Bar chart on the right represent number of cells of each cell type in the single-cell RNA sequencing dataset. (**D** and **E**) Absolute number (**D**) and proportion distribution (**E**) of the 27 cell lineages per sample. Bars represent the absolute number (**D**) or percentage (**E**) of cells assigned to each colour-coded cell lineage relative to the total cells for each sample. (**F**) UMAP illustrating the presence of MAIT cells in both WT and $Mr1^{-/-}$ mice. MAIT cells were identified using TCR sequencing results generated by Cell Ranger. A cell is classified as a MAIT cell if it belongs to a clonotype with expression of Trav1 and Traj33 (TRA), or a clonotype with the expression of Trav1 and Traj33, paired with either Trbv19 or Trbv13 (TRA & TRB) (https://support.10xgenomics.com/single-cell-vdj/software/pipelines/latest/algorithms/inkt-mait).

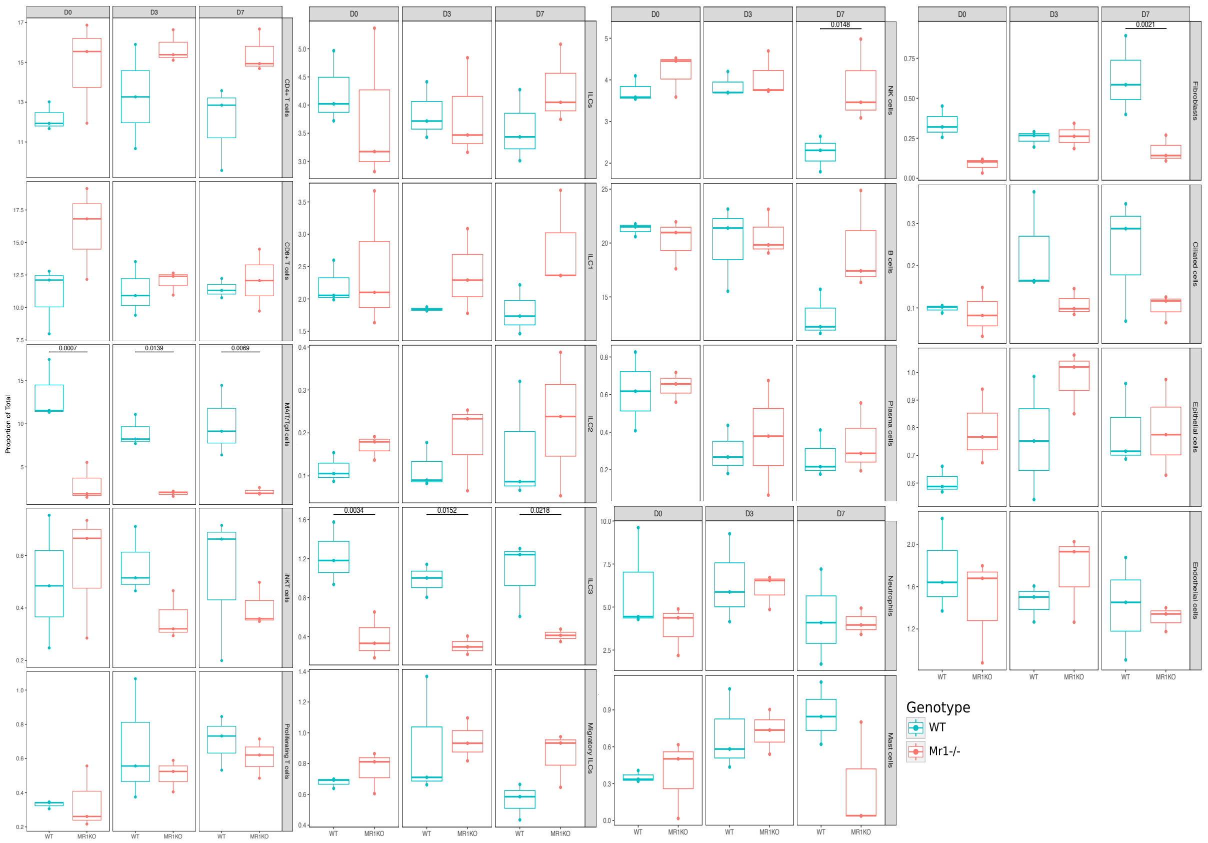


**Supplementary Fig. 8. Cell type distribution in both WT and Mr1$^{-/-}$ mouse lungs post-bleomycin challenge.** Proportion of the indicated cell types of total lung cells was calculated for individual mice at the indicated time points after bleomycin challenge and for PBS treated control mice (n = 3 for each genotype). *P* values generated using a two-way ANOVA with Sidak's multiple comparisons test.

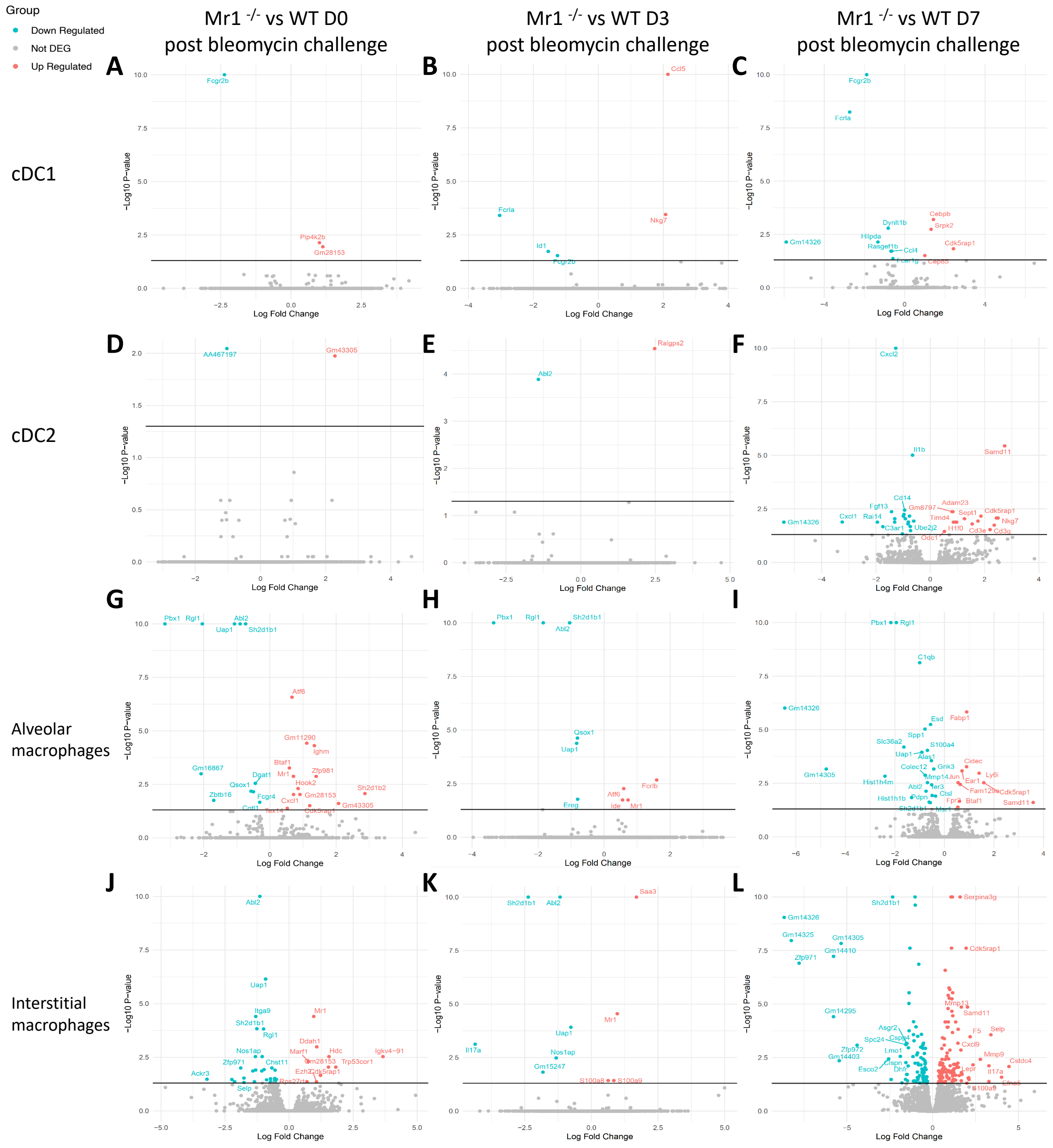


**Supplementary Fig. 9. Differential gene expression in dendritic cell and macrophage populations in Mr1$^{-/-}$ vs. WT mice in response to bleomycin.** Volcano plots showing cDC1 (**A** to **C**), cDC2 (**D** to **F**), alveolar macrophages (**G** to **H**) and interstitial macrophage (**J** to **L**)-specific DEGs based on adjusted *P* value < 0.05 between Mr1$^{-/-}$ and wild-type mice before and post-bleomycin challenge. Horizontal line indicates *P* value threshold of 0.05.

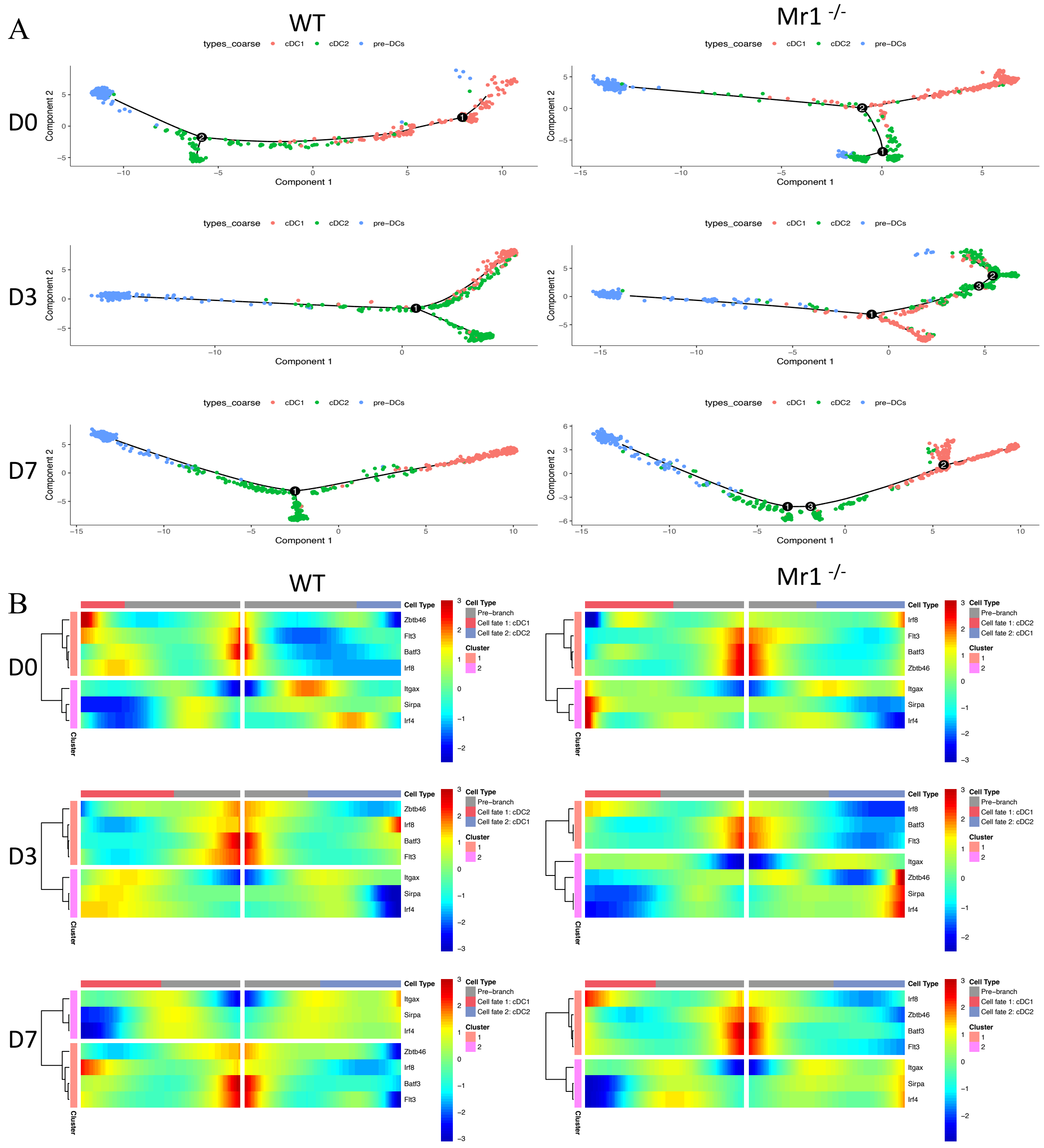


**Supplementary Fig. 10. Differentiation trajectories and gene expression patterns of pre-DCs to cDC1 and cDC2 in WT and Mr1$^{-/-}$ mouse lungs.** (**A**) Analysis using Monocle2 [2] showing branches of differentiation trajectory from pre-DCs, the progenitors of cDCs, towards cDC1 and cDC2. (**B**) Heatmap showing dynamic change in gene expression of two differentiation branches (cDC1 and cDC2). Higher gene expression levels are denoted in red, while lower levels in blue.

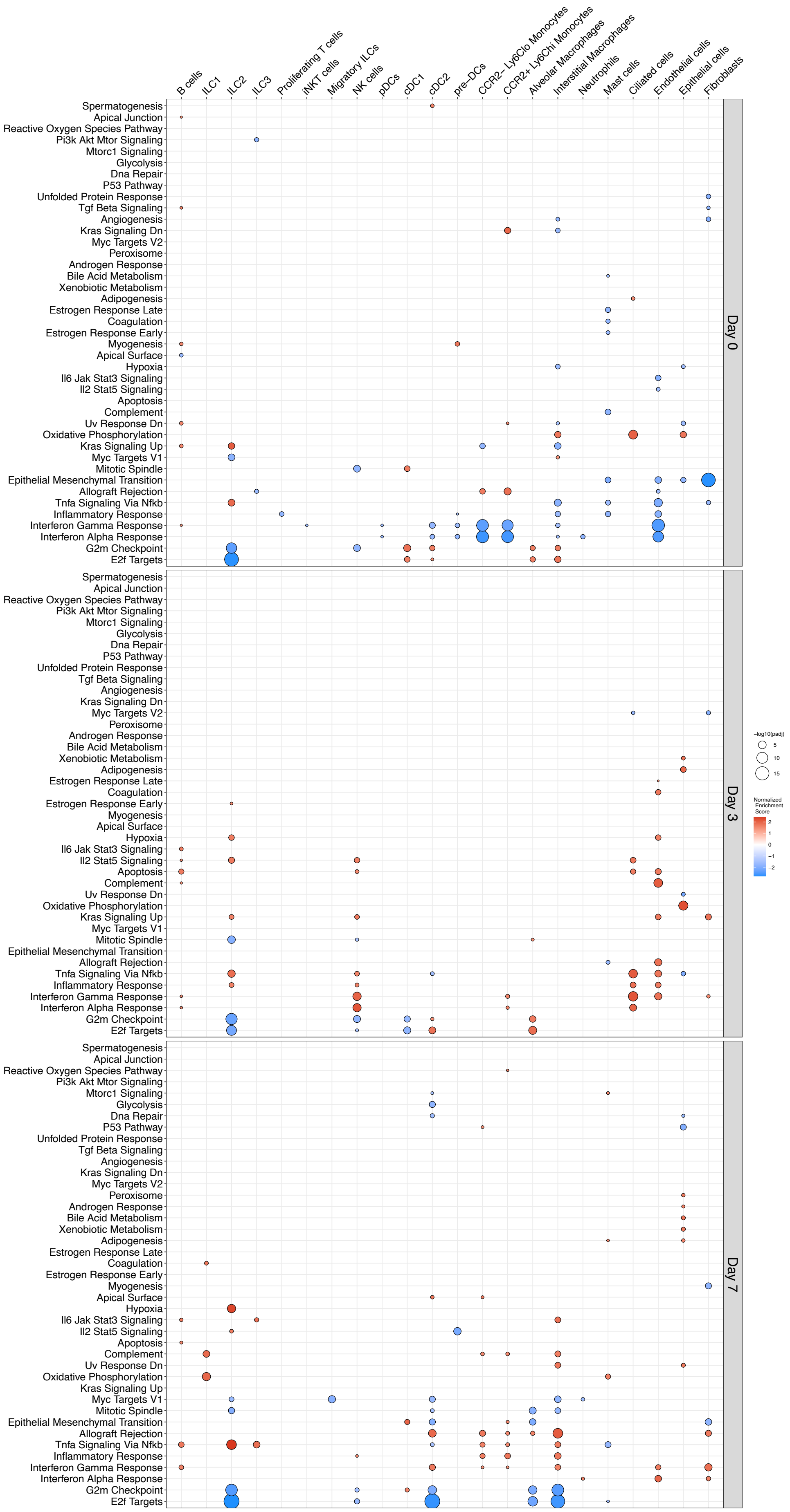
B cells
ILC1
ILC2
ILC3
Proliferating T cells
iNKT cells
Migratory ILCs
NK cells
pDCs
cDC1
cDC2
pre–DCs
CCR2– Ly6Clo Monocytes
CCR2+ Ly6Chi Monocytes
Alveolar Macrophages
Interstitial Macrophages
Neutrophils
Mast cells
Ciliated cells
Endothelial cells
Epithelial cells
Fibroblasts
Spermatogenesis
Apical Junction
Reactive Oxygen Species Pathway
Pi3k Akt Mtor Signaling
Mtorc1 Signaling
Glycolysis
Dna Repair
P53 Pathway
Unfolded Protein Response
Tgf Beta Signaling
Angiogenesis
Kras Signaling Dn
Myc Targets V2
Peroxisome
Androgen Response
Bile Acid Metabolism
Xenobiotic Metabolism
Adipogenesis
Estrogen Response Late
Coagulation
Estrogen Response Early
Myogenesis
Apical Surface
Hypoxia
Il6 Jak Stat3 Signaling
Il2 Stat5 Signaling
Apoptosis
Complement
Uv Response Dn
Oxidative Phosphorylation
Kras Signaling Up
Myc Targets V1
Mitotic Spindle
Epithelial Mesenchymal Transition
Allograft Rejection
Tnfa Signaling Via Nfkb
Inflammatory Response
Interferon Gamma Response
Interferon Alpha Response
G2m Checkpoint
E2f Targets
Day 0
Day 3
Day 7
−log10(padj)
5
10
15
Normalized Enrichment Score
2
1
0
−1
−2

**Supplementary Fig. 11. Gene Set Enrichment Analysis in lung cells of Mr1$^{-/-}$ and WT mice post-bleomycin challenge.** Gene Set Enrichment Analysis (GSEA) performed on the Mr1$^{-/-}$ and wild-type (WT) mice across different time points utilizing the MSigDB Hallmark collection. Log2 fold changes in differential expression, as computed through DESeq2, served as the rank statistics. The color of the dots represents the Normalised Enrichment Score (NES), while their size indicates the -log10-adjusted *P* value. *P* value was modified using the Benjamini-Hochberg method. The direction of the NES signifies the relative increase or decrease in the Mr1$^{-/-}$ mice when compared to the WT mice.

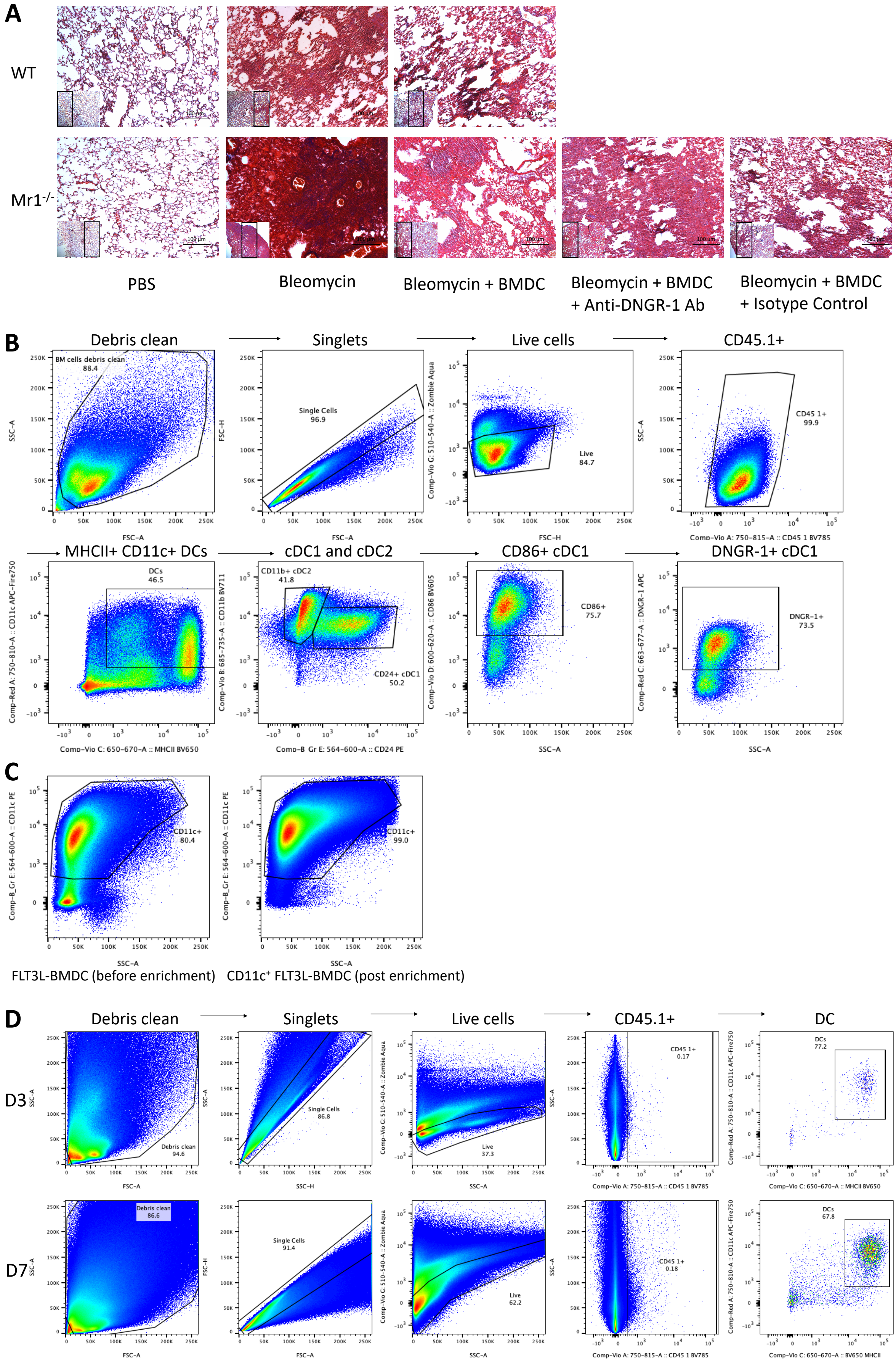
A
WT
Mr1-/-
PBS
Bleomycin
Bleomycin + BMDC
Bleomycin + BMDC + Anti-DNGR-1 Ab
Bleomycin + BMDC + Isotype Control
B
Debris clean
Singlets
Live cells
CD45.1+
BM cells debris clean 88.4
Single Cells 96.9
Live 84.7
CD45 1+ 99.9
MHCII+ CD11c+ DCs
cDC1 and cDC2
CD86+ cDC1
DNGR-1+ cDC1
DCs 46.5
CD11b+ cDC2 41.8
CD24+ cDC1 50.2
CD86+ 75.7
DNGR-1+ 73.5
C
CD11c+ 80.4
CD11c+ 99.0
FLT3L-BMDC (before enrichment)
CD11c+ FLT3L-BMDC (post enrichment)
D
Debris clean
Singlets
Live cells
CD45.1+
DC
D3
D7
Debris clean 94.6
Single Cells 86.8
Live 37.3
CD45 1+ 0.17
DCs 77.2
Debris clean 86.6
Single Cells 91.4
Live 62.2
CD45 1+ 0.18
DCs 67.8

**Supplementary Fig. 12. Lung tissue response and dendritic cell characterisation in WT and Mr1$^{-/-}$ mice post-bleomycin challenge and -adoptive transfer of FLT3L-BMDC.** (**A**) Representative images of lung slices of PBS or bleomycin-challenged WT and Mr1$^{-/-}$ mice at D21, stained with Masson's trichrome. (**B**) Gating scheme for the identification of conventional dendritic cells (cDC) in Fms-like tyrosine kinase-3 ligand (Flt3L)-generated bone marrow-derived dendritic cells (FLT3L-BMDC), and expression of CD86 and DNGR-1 on cDC1. cDCs are identified as CD11c$^{+}$ MHCII$^{+}$ cells and can be divided into cDC1 (CD24$^{+}$ CD11b$^{-}$) and cDC2 (CD24$^{-}$ CD11b$^{+}$). (**C**) Representative flow plot showing the CD11c$^{+}$ FLT3L-BMDC population before and post-CD11c enrichment (gated on all live CD45.1$^{+}$ cells). (**D**) Flow cytometric analysis of adoptive-transferred CD11c$^{+}$ bone marrow-derived dendritic cells (BMDC, gated as CD45.1$^{+}$ MHCII$^{+}$ CD11c$^{+}$ population) in Mr1$^{-/-}$ mice lungs at day 3 and day 7 post-adoptive transfer, respectively.

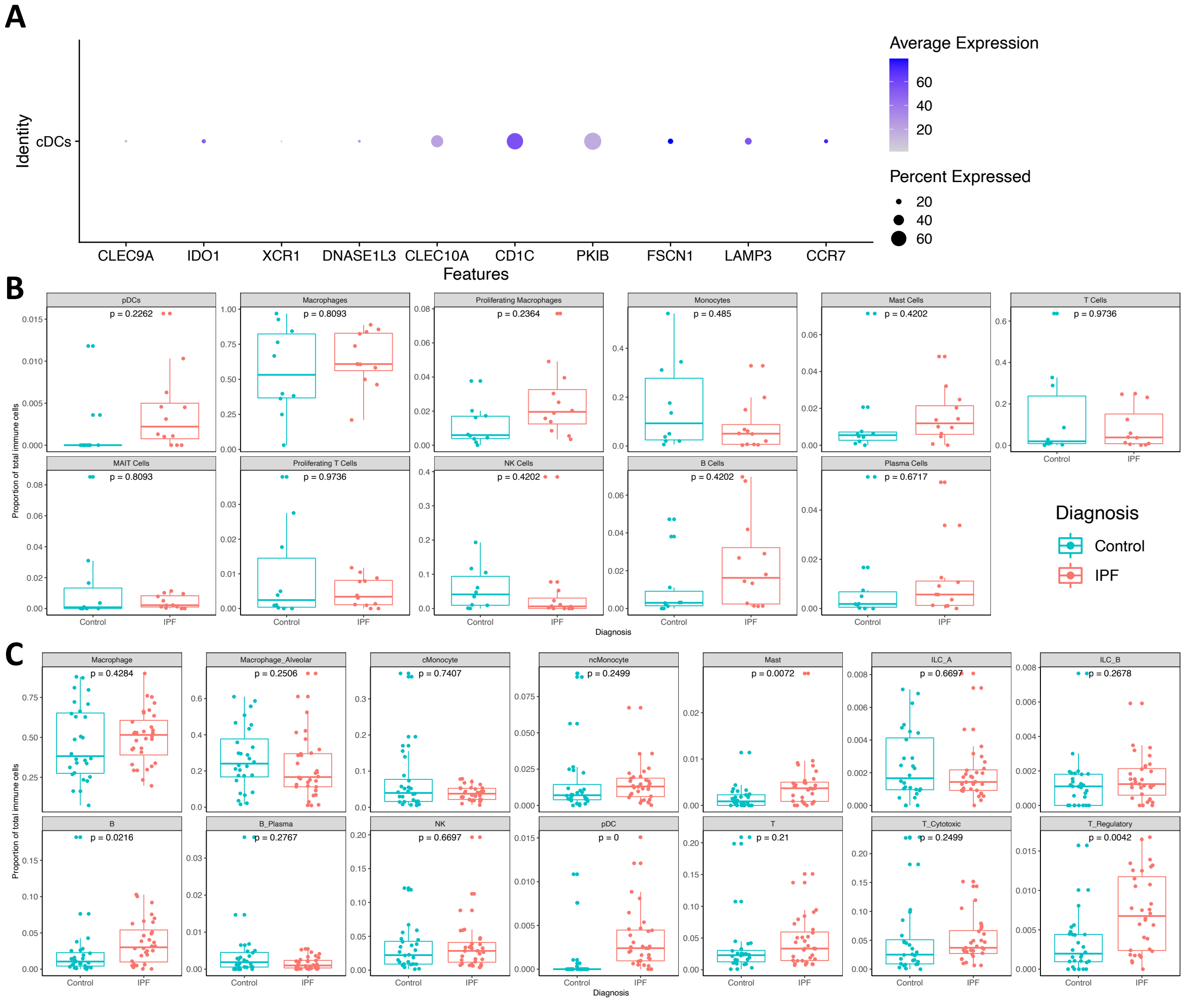


**Supplementary Fig. 13. Gene expression profiling of cDCs in IPF patients' lungs and comparative analysis of cell frequencies between IPF patients and controls.** (**A**) Dot plot showing the gene expression profile for the cDCs in the GSE135893 dataset. Dot size represents percentage of expressed cells. Colour scales denote the average gene expression. (**B** and **C**) Boxplots present frequencies of indicated cell type, as proportions of total lung cells, in IPF patients versus controls, sourced from GSE135893 (**B**) and GSE136831 (**C**). *P* values generated using a two-way ANOVA with Sidak's multiple comparisons test.

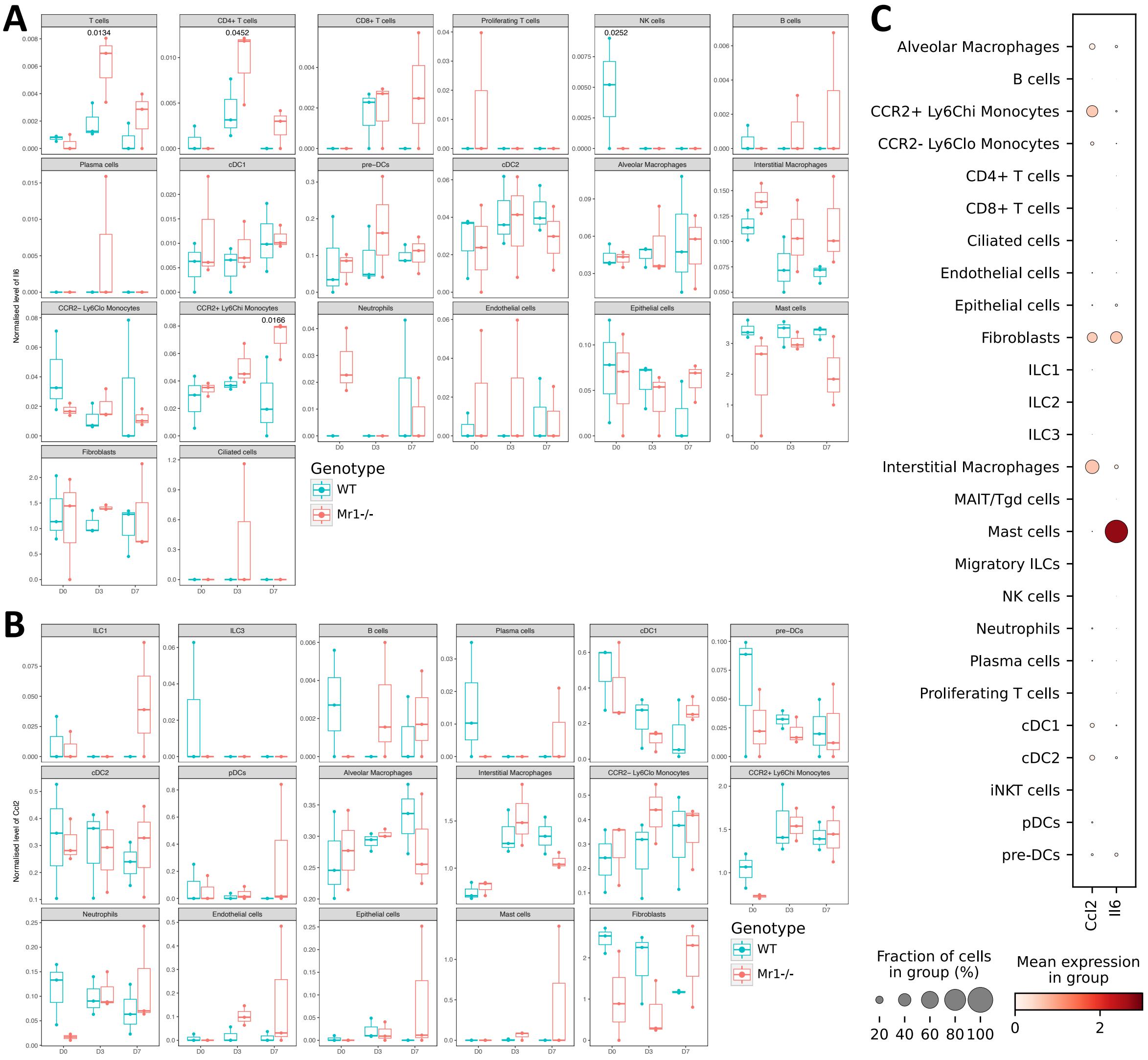


**Supplementary Fig. 14. Differential expression of inflammatory markers Il6 and Ccl2 in lung cell types pre- and post-bleomycin challenge.** Bar plots showing the gene expression of Il6 (**A**) and Ccl2 (**B**) across various cell types pre and post-bleomycin challenge. (**C**) Dot plot illustrating the combined gene expression levels of Il6 and Ccl2 across different cell types, collated from all time points and both WT and Mr1$^{-/-}$ mice.

Supplementary Table 1. DEGs of MAIT cells, IPF vs Control

| | **Log2 Fold Change** | ***P* value** | **Adjusted *P* value** |
|---|---|---|---|
| CTD-3252C9.4 | 3.40847694 | 5.06E-09 | 0.00013339 |
| TNFSF9 | 3.00877575 | 1.20E-07 | 0.00316397 |
| CCL4L2 | 2.67002871 | 1.65E-10 | 4.36E-06 |
| ITM2C | 2.39843978 | 2.32E-09 | 6.13E-05 |
| HSPA1B | 2.37900959 | 1.22E-20 | 3.21E-16 |
| DUSP4 | 2.37889768 | 7.97E-08 | 0.0021023 |
| CDKN1A | 2.20750059 | 7.27E-07 | 0.01917007 |
| HSPA1A | 2.01768082 | 5.54E-17 | 1.46E-12 |
| RGS1 | 1.95067011 | 4.28E-16 | 1.13E-11 |
| PHLDA1 | 1.91687999 | 4.00E-10 | 1.06E-05 |
| CCL3 | 1.86015604 | 1.26E-07 | 0.00332594 |
| NR4A1 | 1.82189329 | 2.85E-10 | 7.51E-06 |
| EGR1 | 1.78128101 | 5.63E-08 | 0.00148433 |
| CSRNP1 | 1.74935216 | 9.58E-07 | 0.02526935 |
| SCGB3A1 | 1.72801781 | 3.13E-17 | 8.25E-13 |
| SCGB1A1 | 1.69915943 | 9.92E-14 | 2.62E-09 |
| SERTAD1 | 1.69208413 | 6.91E-07 | 0.01821557 |
| PDE4B | 1.64592784 | 1.05E-07 | 0.00275658 |
| BIRC3 | 1.61577906 | 1.66E-07 | 0.00437926 |
| FOSB | 1.50301559 | 4.32E-10 | 1.14E-05 |
| DNAJB1 | 1.24869004 | 8.04E-15 | 2.12E-10 |
| DNAJA1 | 1.22687395 | 3.24E-12 | 8.53E-08 |

| | | | |
|---|---|---|---|
| RGCC | 1.21337533 | 5.22E-09 | 0.00013778 |
| RPS4Y1 | 1.20033202 | 1.36E-08 | 0.00035806 |
| CORO1B | 1.19148334 | 6.53E-09 | 0.00017236 |
| BTG2 | 1.11280058 | 1.02E-10 | 2.68E-06 |
| CAPG | 1.08636027 | 9.99E-07 | 0.02635963 |
| NR4A2 | 1.07056276 | 6.34E-07 | 0.01673196 |
| CCL4 | 0.98464434 | 2.16E-07 | 0.00569644 |
| FOS | 0.92669826 | 2.02E-17 | 5.32E-13 |
| PPP1R15A | 0.89848942 | 7.88E-07 | 0.02077662 |
| TNFAIP3 | 0.89036899 | 6.43E-10 | 1.70E-05 |
| PTPRCAP | 0.80498537 | 7.99E-10 | 2.11E-05 |
| NFKBIA | 0.69743629 | 1.23E-10 | 3.24E-06 |
| LGALS1 | 0.64854233 | 2.07E-07 | 0.00547243 |
| KLF6 | 0.64467035 | 2.59E-11 | 6.84E-07 |
| ALOX5AP | 0.60331068 | 3.17E-07 | 0.00837339 |
| ZFP36 | 0.58343053 | 1.52E-12 | 4.01E-08 |
| HLA-DRB1 | 0.56565865 | 1.16E-07 | 0.00305218 |
| DYNLL1 | 0.56083732 | 9.13E-07 | 0.02409233 |
| CD69 | 0.54163104 | 3.29E-07 | 0.00868113 |
| DUSP1 | 0.52541753 | 1.67E-10 | 4.41E-06 |
| JUNB | 0.46129731 | 3.00E-08 | 0.00079207 |
| HSP90AA1 | 0.45594655 | 8.42E-11 | 2.22E-06 |
| CRIP1 | 0.42043194 | 6.61E-07 | 0.01744826 |
| S100A11 | 0.41303276 | 6.73E-07 | 0.01775344 |
| CD74 | 0.3775569 | 5.61E-08 | 0.00148106 |

| | | | |
|---|---|---|---|
| FTH1 | 0.34485861 | 1.08E-12 | 2.84E-08 |
| VIM | 0.27986256 | 1.34E-06 | 0.03524605 |
| RPS26 | -0.4863287 | 5.11E-15 | 1.35E-10 |
| PCBP2 | -0.6295141 | 3.05E-16 | 8.04E-12 |
| PRF1 | -0.8782519 | 3.43E-09 | 9.05E-05 |
| TRGV10 | -1.9454736 | 2.45E-08 | 0.00064673 |
| MTRNR2L12 | -2.71823 | 2.10E-07 | 0.00553425 |
| TRAV12-1 | -4.0025993 | 2.32E-09 | 6.13E-05 |

Supplementary Table 2. Flow cytometry reagents

| No. | Antigen | Conjugation | Clone | Host | Reactivity | Isotype | Concentration Titration | Catalogue number | Supplier |
|---|---|---|---|---|---|---|---|---|---|
| 1 | CD103 | AF700 | 2 E7 | Hamster | Mouse | Armenian Hamster IgG | 1:100 | 121442 | BioLegend |
| 2 | CD45 | BUV395 | 30-F11 | Rat | Mouse | Rat IgG2b, κ | 1:100 | 564279 | BD Biosciences |
| 3 | CD11c | APC/Fire™ 750 | N418 | Rat | Mouse | Armenian Hamster IgG | 1:100 | 117352 | Biolegend |
| 4 | CD19 | BV480 | 1D3 | Rat | Mouse | Lewis IgG2a, κ | 1:100 | 566107 | BD Biosciences |
| 5 | MERTK (Mer) | PE-Cy7 | 2B10C42 | Rat | Mouse | Rat IgG2a, κ | 1:100 | 151522 | Biolegend |
| 6 | CD4 | AF488 | GK1.5 | Rat | Mouse | Rat IgG2b, κ | 1:100 | 100423 | Biolegend |
| 7 | Ly-6G | BUV615 | 1A8 | Rat | Mouse | Lewis IgG2a, κ | 1:100 | 751263 | BD Biosciences |
| 8 | Ly-6C | BV570 | HK1.4 | Rat | Mouse | Rat IgG2c, κ | 1:100 | 128029 | Biolegend |
| 9 | MHC II | BV650 | M5/114.15.2 | Rat | Mouse | Rat IgG2b, κ | 1:100 | 107641 | Biolegend |
| 10 | CD8a | BV605 | 53-6.7 | Rat | Mouse | Rat IgG2a, κ | 1:100 | 100744 | Biolegend |
| 11 | CD11b | BV711 | M1/70 | Rat | Mouse/Human | Rat IgG2b, κ | 1:100 | 101242 | Biolegend |
| 12 | CD3 | BV785 | 17A2 | Rat | Mouse | Rat IgG2b, κ | 1:100 | 100232 | Biolegend |

| | | | | | | | | | |
|---|---|---|---|---|---|---|---|---|---|
| 13 | CD64 | PE | X54-5/7.1 | Mouse | Mouse | Mouse IgG1, κ | 1:100 | 139304 | Biolegend |
| 14 | NK1.1 | PE/Cyanine5 | PK136 | Mouse | Mouse | Mouse IgG2a, κ | 1:100 | 108716 | Biolegend |
| 15 | CD44 | BUV805 | IM7 | Rat | Mouse | Rat IgG2b, κ | 1:100 | 741921 | BD Biosciences |
| 16 | TCR γ/δ | BUV737 | GL3 | Hamster | Mouse | Armenian Hamster IgG2, κ | 1:100 | 748991 | Biolegend |
| 17 | CD19 | PerCP Cy5.5 | 6D5 | Rat | Mouse | Rat IgG2a, κ | 1:100 | 115534 | BioLegend |
| 18 | CD69 | FITC | H1.2F3 | Hamster | Mouse | Armenian Hamster IgG1, λ3 | 1:100 | 553236 | BD Biosciences |
| 19 | TCRβ | PE-Cy7 | H57-597 | Hamster | Mouse | Armenian Hamster IgG2, λ1 | 1:100 | 560729 | BD Biosciences |
| 20 | CD25 | APC | PC61.5 | Rat | Mouse | Rat IgG1, λ | 1:100 | 17-0251-82 | eBioscience™ |
| 21 | CD45.2 | BV711 | 104 | Mouse | Mouse | Mouse SJL IgG2a, κ | 1:100 | 563685 | BD Biosciences |
| 22 | IFN-γ | BV650 | XMG1.2 | Rat | Mouse | Rat IgG1, κ | 1:100 | 563854 | BD Biosciences |
| 23 | IL-22 | APC | Poly5164 | Goat | Mouse | Polyclonal | 1:100 | 516409 | BioLegend |
| 24 | IL-10 | APC-Cy7 | JES5-16E3 | Rat | Mouse | Rat IgG2b, κ | 1:100 | 505010 | BioLegend |
| 25 | IL-17A | PE | TC11-18H10.1 | Rat | Mouse | Rat IgG1, κ | 1:100 | 506904 | BioLegend |

| | | | | | | | | | |
|---|---|---|---|---|---|---|---|---|---|
| 26 | GM-CSF | PE/Dazzle™ 594 | MP1-22E9 | Rat | Mouse | Rat IgG2a, κ | 1:100 | 505422 | BioLegend |
| 27 | SiglecH | PerCPCy5.5 | 551 | Rat | Mouse | Rat IgG1, κ | 1:100 | 129614 | BioLegend |
| 28 | CD45 | FITC | 30-F11 | Rat | Mouse | Rat IgG2b, κ | 1:100 | 103108 | BioLegend |
| 29 | CD86 | BV605 | GL-1 | Rat | Mouse | Rat IgG2a, κ | 1:100 | 105037 | BioLegend |
| 30 | CD40 | PE/Cyanine5 | 44986 | Rat | Mouse | Rat IgG2a, κ | 1:100 | 124618 | BioLegend |
| 31 | CD192 | PE/Dazzle™ 594 | SA203G11 | Rat | Mouse | Rat IgG2b, κ | 1:100 | 150636 | BioLegend |
| 32 | CD192 | APC/Fire™ 750 | SA203G11 | Rat | Mouse | Rat IgG2b, κ | 1:100 | 150630 | BioLegend |
| 33 | CD317 | PE/Cyanine7 | eBio927 | Rat | Mouse | Rat IgG2b, κ | 1:100 | 25-3172-82 | eBioscience™ |
| 34 | CD370 | APC | 7H11 | Rat | Mouse | Rat IgG1, κ | 1:100 | 143506 | BioLegend |
| 35 | CD24 | PE/Dazzle™ 594 | M1/69 | Rat | Mouse | Rat IgG2b, κ | 1:100 | 101838 | BioLegend |
| 36 | CD45.2 | FITC | 104 | Mouse | Mouse | Mouse (SJL) IgG2a, κ | 1:100 | 109806 | BioLegend |
| 37 | CD45.1 | BV785 | A20 | Mouse | Mouse | Mouse (A.SW) IgG2a, κ | 1:100 | 110743 | BioLegend |
| 38 | CD11c | PE | N418 | Hamster | Mouse | Armenian Hamster IgG | 1:100 | 117308 | BioLegend |

| | | | | | | | | | |
|---|---|---|---|---|---|---|---|---|---|
| 39 | CD45R | BV421 | RA3-6B2 | Rat | Mouse | Rat IgG2a, κ | 1:100 | 103251 | BioLegend |
| 40 | Zombie NIR™ Fixable Viability Kit | | | | | | 1:1000 | 423106 | BioLegend |
| 41 | Zombie Aqua™ Fixable Viability Kit | | | | | | 1:1000 | 423102 | BioLegend |
| 42 | Zombie Yellow™ Fixable Viability Kit | | | | | | 1:1000 | 423104 | BioLegend |

**Supplementary Data 1. DEGs of MAIT cells, WT B6 bleomycin challenged versus WT B6 unchallenged (separate file)**

Differentially expressed genes [log2 fold change (FC) > 1, adjusted $P < 0.05$] of mouse lung MAIT cells on days 3, 7, 14, 21 and 28 post bleomycin challenge, respectively, compared with unchallenged PBS controls.

**Supplementary Data 2. DEGs of MAIT cells, WT B6 PR8 virus infected versus WT B6 uninfected (separate file)**

Differentially expressed genes [log2 fold change (FC) > 1, adjusted $P < 0.05$] of mouse lung MAIT cells on days 3, 7, 14, 21 and 28 post PR8 influenza A virus infection, respectively, compared with uninfected PBS controls.

**Supplementary Data 3. DEGs of total lung, Mr1$^{-/-}$ versus WT B6 (separate file)**

Differentially expressed genes [log2 fold change (FC) > 1, adjusted $P < 0.05$] in whole lung tissue between Mr1$^{-/-}$ and WT mice lungs on days 0, 3, 7, 14 and 21 post bleomycin challenge, respectively.

**Supplementary Data 4. DEGs, cell type-specific, WT B6 versus Mr1$^{-/-}$ (separate file)**

Differences in cell type-specific differentially expressed genes between WT and Mr1$^{-/-}$ mice on days 0 (unchallenged PBS controls), 3 and 7 post bleomycin challenge, respectively.
